# Variability in intrinsic drug tolerance in *Mycobacterium tuberculosis* corresponds with phylogenetic lineage

**DOI:** 10.1101/2025.07.04.663131

**Authors:** Valerie F. A. March, Michaela Zwyer, Chloé Loiseau, Daniela Brites, Galo A. Goig, Selim Bouaouina, Anna Doetsch, Miriam Reinhardt, Sevda Kalkan, Sebastien Gagneux, Sonia Borrell

## Abstract

Drug tolerance allows bacteria to survive extended exposure to bactericidal drugs and is thought to play a role in drug resistance evolution. In *Mycobacterium tuberculosis* (Mtb), the causative agent of tuberculosis (TB), multidrug resistant TB (MDR-TB) outbreaks are frequently caused by strains belonging to two phylogenetic lineages of the human-adapted strains of the Mtb Complex, namely lineages (L) 2 and L4. We hypothesized that members of L2 and L4 are more intrinsically drug tolerant, and as such, more readily evolve drug resistance. To explore this, we devised a high throughput *in vitro* assay to measure drug tolerance in Mtb. We selected a cohort of strains representative of the globally most frequent lineages L1 – L4. We measured tolerance to rifampicin and bedaquiline and found L3 and L4 strains to have higher tolerance compared to L1 and L2 strains. In addition, phylogenetically closely related strains exhibited similar levels of tolerance, suggesting that tolerance is heritable. Finally, we explored genes previously reported to be associated with tolerance in Mtb and found significant enrichment in mutations in genes involved in cell wall and cell processes, intermediary metabolism and respiration, as well as lipid metabolism in high-tolerance strains.

## Introduction

Antibiotics are key in the global fight against tuberculosis (TB), which remains the most frequent cause of human death due to a single infectious agent (1). Human TB is mainly caused by *Mycobacterium tuberculosis* (Mtb). With a lipid rich cell wall and the ability to go into a dormant non-replicative state (2, 3), Mtb is canonically understood to be intrinsically tolerant to many antibiotics. While this is true, there are effective antibiotics to treat TB, where efficacy is on average 88% for drug susceptible cases (1).

Drug-resistant TB has historically been challenging to treat, with recent reports showing efficacy around 65% (1). Multidrug-resistant TB (MDR-TB) is defined by resistance to the two key frontline drugs rifampicin (RIF) and isoniazid (INH), which are the main components of susceptible TB treatment(4). Patients receive RIF and INH for up to 6-months, with the first two months being an intensive phase including pyrazinamide and ethambutol (4). Treatment of MDR-TB has recently been updated with the introduction of short all-oral regimens containing the novel anti-TB drugs bedaquiline (BDQ) and pretomanid, alongside linezolid and moxifloxacin where appropriate (5). While drug resistance affects ∼3% of new cases globally, it remains a barrier for TB eradication (1). Moreover, in geographical hot spots of MDR-TB, up to 54% of the disease burden can be caused by drug-resistant strains (1), which also have the ability to transmit from patient to patient (6–9). Furthermore, the threat of resistance to the new drugs remains. As such, there is a need to better understand how drug resistance emerges.

Globally, MDR-TB outbreaks tend to be caused by strains of certain phylogenetic lineages (10). Mtb exists within a larger family termed the *Mycobacterium tuberculosis* complex (MTBC) containing both human and animal-adapted strains (11). Human-adapted strains comprise 10 lineages. Globally, lineages (L) 1 – 4 are responsible for the majority of the TB burden (11, 12). MDR-TB is frequently attributed to strains belonging to L2 and L4 (7, 8, 13), but the basis of this association is not fully understood.

Recent literature has given a role to drug tolerance in drug resistance evolution (14–16). Drug tolerance is a phenotype wherein bacterial populations can withstand extended durations at killing concentrations of bactericidal antibiotics (17). In fast growing bacteria, this has been linked to resistance emergence in patients, with tolerance being an intermediate phenotype on the path to resistance (14). Drug tolerance is thus an important phenotypic characteristic in drug-pathogen interaction; however, the role of drug tolerance in Mtb drug resistance evolution remains to be established.

We hypothesized that MTBC lineages show differences in intrinsic drug tolerance, which could account for lineage patterned propensities for drug resistance evolution. Specifically, we asked whether strains of L2 and L4 are more intrinsically drug tolerant and thus more readily acquire drug resistance. To address this question, we adapted a high throughput *in vitro* assay to measure drug tolerance in a diverse set of Mtb strains across L1 – L4 and explored genetic polymorphisms associated with drug tolerance in Mtb.

### Methods

Strain selection

We selected strains from our BSL3 biobank genomically predicted to be drug susceptible according to the WHO clinical definition, and members of MTBC lineages L1 –L4. For L1 and L3, we selected strains related to the three most globally prevalent sublineages, and for L2 and L4 we selected two globally prevalent sublineages and one geographically restricted sublineage (see supplementary data). Strains were collected from different global sites as such our final strain set included phylogenetically and geographically diverse strains (Detailed in **Table S2**). Within this strain panel, we included a previously published reference clinical strains (18) set.

### Strain culture and stock preparation

Strains were inoculated from the biobank into 10 ml Middlebrook 7H9 liquid medium supplemented with 10% ADC (7H9 ADC, 5% bovine albumin-fraction V + 2% dextrose + 0.003% catalase, Sigma-Aldrich), 0.5% glycerol (PanReac AppliedChem) and 0.1% Tween-80 (Sigma-Aldrich), which will henceforth be referred to as 7H9 ADC. Cultures were grown until early stationary phase (OD_600nm_ 0.8 – 1) at 37°C shaking at 140 rpm, and then centrifuged at 1000 *x g*, after which the resulting supernatant was discarded. Pellets were resuspended in 2 ml 7H9 ADC + 5% glycerol and aliquoted to into working stocks in 250 µl volumes. Working stocks were frozen at - 80 °C.

In order to work with strains in parallel, calibrated stocks were prepared. To do so, 50 µl of working stocks were inoculated into 10 ml 7H9 ADC, which were left to grow until mid-log phase (OD_600nm_ 0.5 – 0.7). Once turbid, cultures were centrifuged at 260 *x g* to pellet clumps. 9 ml of upper culture supernatant was transferred into a clean tube, and the OD_600nm_ measured. Stocks were calibrated into 10 ml cultures at OD_600nm_ 0.1 in 7H9 ADC, aliquoted into 1 ml calibrated stocks, and frozen at - 80 °C. For use, calibrated stocks were thawed to at ambient temperature, any remaining culture after use was discarded.

### Preparation of antibiotic stocks

Antibiotic stocks were prepared from powdered stocks of rifampicin (RIF, Sigma-Aldrich), bedaquiline fumarate (BDQ, MedChem Express), moxifloxacin (Mfx, Sigma) and Pretomanid (Ptm, Sigma). Working stocks were diluted in dimethyl sulfoxide (DMSO, AppliChem).

### Preparation of resazurin stocks

Resazurin was prepared by dissolving 3.125 mg of resazurin sodium salt (Sigma-Aldrich) in 25 ml of Milli-Q water. The resazurin solution was vortexed and filter sterilized using a 0.22 μM syringe filter in a sterile biosafety cabinet. Aliquots of 1ml were subsequently frozen. Fresh aliquots were thawed when needed and any excess was discarded after each use.

### Determining strain minimum inhibitory concentration (MIC)

Strain MICs were determining by the REMA method as previously described (19–21). In short, antibiotic plates were prepared using a 2-fold dilution series in triplicate. RIF and BDQ series were prepared on a single plate and 10 µl per well of a single strain was inoculated per plate, with drug free wells included for positive controls and drug free culture-free wells included for negative controls. Plates were incubated at 37°C with 5% CO_2_ for 7-9 days. Differences in incubation time were due to different strain growth propensities. After incubation, 10 µl of resazurin was inoculated into each well, after which plates were incubated for a further 24 hours. Fluorescence was measured using the Tecan Infinite Pro 200, with excitation at 560 nm and emission measured at 590 nm. Relative growth of each well was calculated by subtracting the average background fluorescence of negative controls well and expressing growth relative to the average signal in drug-free positive control wells. The MIC was defined as the drug concentration at which growth was inhibited by at least 90%. The MIC assay was performed twice before the MIC was set for each strain.

### BacTiter-Glo linearity test

To assess whether ATP signal measured with the BacTiter-Glo assay changes in a linear fashion according to bacterial density, we tested linearity in a reference set of strains from L1, L2, L3 and L4. Based on the premise that ATP signal should change in a manner proportional to bacterial density. We used three representative strains of each lineage from the reference set as previously published (18, 22). Calibrated strain stocks were thawed and inoculated at a 50 µl volume in 950 µl of 7H9 ADC in triplicate. A ten-fold dilution was performed twice to create a series of 2-log difference. Following this, 100 µl of the resulting suspensions was inoculated into luminescence plates pre-loaded with 100 µl BacTiter-Glo reagent (Promega) prepared according to manufacturer’s instructions. Plates were left to lyse and incubate for 35 minutes and luminescence values were measured using the Tecan Spark multimodal plate reader.

### Reference CFU-based Time-killing assay

To measure tolerance using a gold-standard approach, we used a previously published reference set of clinical strains from L1 – L4(18). We inoculated 50 µl of working stocks into 10 ml 7H9 ADC to generate starter cultures. Once cultures reach mid-log phase they were calibrated to an OD of 0.1 using the method described above. 500 µl of resulting calibrated culture was subsequently inoculated into 7H9 ADC with or without RIF at 25 X MIC to a total volume of 10 ml. Drug-free cultures were sampled for CFU enumeration by inoculating 20 µl of experimental culture in triplicate into dilution plates containing 180 µl of PBST (Phosphate Buffer Saline + 0.05% Tween80, Sigma-Aldrich). Samples were then plated for colony forming unit (CFU) enumeration on BBL^TM^ Middlebrook 7H11 agar (Becton – Dickenson) plates supplemented with oleic acid, albumin, dextrose and catalase (OADC, 0.05% oleic acid (Axonlab) in ADC). Experimental tubes were left to shake at 140 rpm at 37 °C.

Cultures were sampled every second day post inoculation for 8 days. To assess survival 1 ml of culture was sampled and inoculated into a 2 ml screwcap micro centrifuge tube. Samples were washed twice by centrifugation at 7000 rpm in a micro centrifuge and re-suspended with fresh 7H9 ADC, as previously described (23). After washing, resulting cultures were plated for CFU enumeration as above.

Survival per time point was defined as the ratio of CFUs in the drug condition over the initial CFU concentration from the untreated control at experiment initiation. These values were fed into a python script that considers adjacent survival values as coordinate points on a decay function where x is the survival and y is the time (GetMDK, https://git.scicore.unibas.ch/TBRU/mdkcalculator). It searches for the pair that span the time point at which survival would be at a specific threshold, and extrapolates a linear function. It then uses this function to solve for the time at which the survival would be at the given threshold. For our purposes, we used 5% survival, allowing us to calculate the minimum duration for killing 95% of the bacterial population (MDK_95_). We used the function set at log = true.

### A high throughput real-time time killing assay (rt-TKA)

To overcome experimental limitations in handling multiple Mtb strains in parallel, we devised a real-time time killing assay to measure tolerance. The method relies on using frozen strain stocks calibrated to the same initial density, inoculated into antibiotics at 400 X MIC in a 96 deep well plate format. Outer wells of deep well plates were filled with media to create a moisture barrier for long-term incubation. 7H9 ADC was dispensed in triplicate at a volume of 950 µl with and without drug at 400 X MIC of RIF or BDQ for each qualifying strain. The maximum DMSO concentration used was 2%.

BacTiter-Glo plates were prepared in bulk for the entire experiment timespan. BacTiter-Glo reagent was prepared according to the manufacturer’s instructions. Using an Integra 96 channel electronic pipette, white luminescence plates (Greiner) were prepared with 100 µl BacTiter – Glo reagent per well and frozen at – 80 °C until needed. For each time point BacTiter-Glo plates were thawed overnight at 4 °C and equilibrated at ambient temperature before use.

In the BSL3, 50 µl thawed calibrated stocks of strains were inoculated in triplicate in their designated wells. Wells were thoroughly mixed with a multichannel pipette and 100 µl of the resulting culture suspension was transferred to the BacTiter-Glo plate. Three wells were inoculated with 7H9 ADC as a background luminescence control and BacTiter-Glo plates were incubated for 30-35 minutes before luminescence was measured. Luminescence was measured using the Tecan Spark Multimodal plate reader. Negative control luminescence values were subtracted from all measurements and the average luminescence in drug free wells for each strain were used as a baseline value for subsequent killing measurement. Inoculated deep well plates were covered with a lid and placed into medium Ziploc bags inside a container. Plates were incubated at 37°C with 5% CO_2_.

At each time point, wells were thoroughly mixed and a 100 µl culture suspension was transferred to a thawed BacTiter-Glo plate. Plates were incubated and measured as described above. For BDQ, strains were sampled two days after inoculation, and every four days thereafter. For RIF, strains were sampled every four days after inoculation. Plates were sampled for 2 weeks and then a final sample was taken at 21 days post inoculation. Survival was set as the signal at a given time point over the initial luminescence signal. Well data where values were negative after correcting for background luminescence were set to zero. Subsequently MDK_95_ values were calculated for each strain using the GetMDK function.

### Statistical analysis

Experimental data were visualized and analysed using GraphPad Prism (version 8.2.1) and R version 4.3.1. Generally, data was inspected for normality and ANOVAs, or T-tests were performed or non-parametric counterparts where appropriate, specific tests are detailed in figure captions.

### Whole-genome sequence analysis

Strains used in this study were previously sequenced and mapped using our in-house genomics pipeline. Briefly, raw fastq files were first analysed with kraken v.1.1.1 to taxonomically classify sequencing reads and identify reads belonging to the MTBC. Trimmomatic v0.39 was then used to remove Illumina adapters and trim reads with a 5 bp sliding window (cutting off when the median quality dropped below 20), and to discard resulting reads shorter than 20 bp. For paired-end data, reads overlapping at least 15bp were merged using SeqPrep v1.3.1. Reads classified by Kraken as non-MTBC were removed. BWA v0.7.17 was used to align the resulting high-quality MTBC reads to the inferred ancestor of the MTBC (10.5281/zenodo.3497110). Duplicate reads were flagged with the MarkDuplicates module of Picard v2.26.2. The mutect2 module of GATK v4.2.3 was used for variant calling. Variants were then filtered with the FilterMutectCalls in microbial mode. Variant annotation and their effects on genes were predicted using SnpEff v5.0. We removed variants falling in repetitive regions for downstream analysis using published lists (24, 25) and such falling in regions annotated as “PE/PPE/PGRS”, “maturase”, “phage”, or “insertion sequence”. Based on the second version of the WHO catalogue, drug resistance-conferring variants were annotated and only mutations having a frequency >= 10% were kept for downstream analyses.

### Phylogenetic analyses

Phylogenetic trees were constructed from alignments of variable positions with less than 10% missing data. We used RaxML v8.2.11 (26) with the general time reversible model of sequence evolution (-m GTRCAT -V), a rapid bootstrap analysis with 100 bootstraps and search for the best-scoring maximum-likelihood phylogeny. Additionally, we adjusted for the fact that only variable positions were included by ascertainment bias correction by Lewis(27) (--asc-corr=lewis). The phylogenetic trees were rooted using a *M. canettii* genome as outgroup (SAMEA2067187).

### Heritability Analysis

To investigate whether closely related strains exhibit similar phenotypes with respect to MIC and BDQ, we used phylosig function from the phytools (28) package. We computed phylogenetic signals with Pagel’s Lambda (29) and Blomberg’s K (30) in our strain set and assessed significance with 1000 simulations tested per phenotype. Detailed insights into the method can be found in the listed references. Briefly, a maximum likelihood tree and continuous phenotypic data are used to test whether data correspond to a Brownian Motion (BM) evolutionary model. When lambda and K =1, closely related organisms resemble each other to the degree one would expect following a BM model. At lambda and K > 1 the relatives resemble each other more than expected, and at lambda and K < 1 the resemblance is intermediate or lower than expected. At zero relatives, do not resemble each other. The function also tests whether these computed phylogenetic signals are statistically significant.

### GWAS on drug tolerance

A GWAS was performed for BDQ and RIF, respectively, on 95 genomes with experimental phenotypic drug tolerance data using Gemma v0.98.1. Fixed (>=90% allele frequency) indels and SNPs were investigated. PLINK v2.00a2.3 was used to filter for variants with an allele frequency of at least 5% resulting in 2317 variants thereof 2033 being analysed by GEMMA. A significance threshold of 2.46 e-5 was chosen based on the 2164 variants that were analysed (0.05/2033). The borderline significant positions for BDQ and the most significant for RIF were visualized on a phylogenetic tree using the ggtree package from R (31).

### Assessing enrichment for mutations in tolerance-associated genes

We compiled a list of genes associated with drug tolerance or persistence in Mtb from various sources (specified in supplementary information). From strain sequences we created a matrix of every mutation detected in the genes list whether they we were present in each strain. We compiled two lists of upper quartile strains from the BDQ and RIF experiments to investigate enrichment in mutations. Using Fisher’s Exact test, we analysed which mutations were significantly enriched in upper quartile strains compared to the rest of the strain set for each experiment. We used Benjamini-Hochberg’s correction with q set to 0.01 to control for false discoveries at a rate of 1%, significant enrichment was defined at p < 0.05.

## Results

### Establishing a high-throughput time killing assay to measure drug tolerance

We set up a real-time time killing assay (rt-TKA) to measure drug tolerance in Mtb. Survival was assessed by measuring intracellular ATP using the BacTiter-Glo assay (**Methods, Figure 1**). ATP signal measured with the BacTiter-Glo reagent has previously been established to correlate with viable bacterial density (Ref). We confirmed that this correlation holds irrespective of Mtb phylogenetic lineage (**Figure 2A**). An initial pilot of the method using lab strain H37Rv was performed using several anti-TB drugs and proved to be reproducible, especially in RIF and BDQ, with similar MDK_95_ values derived from different sampling time-point regimes (**Figure 2B, Figure S1**).

**Figure 1.**
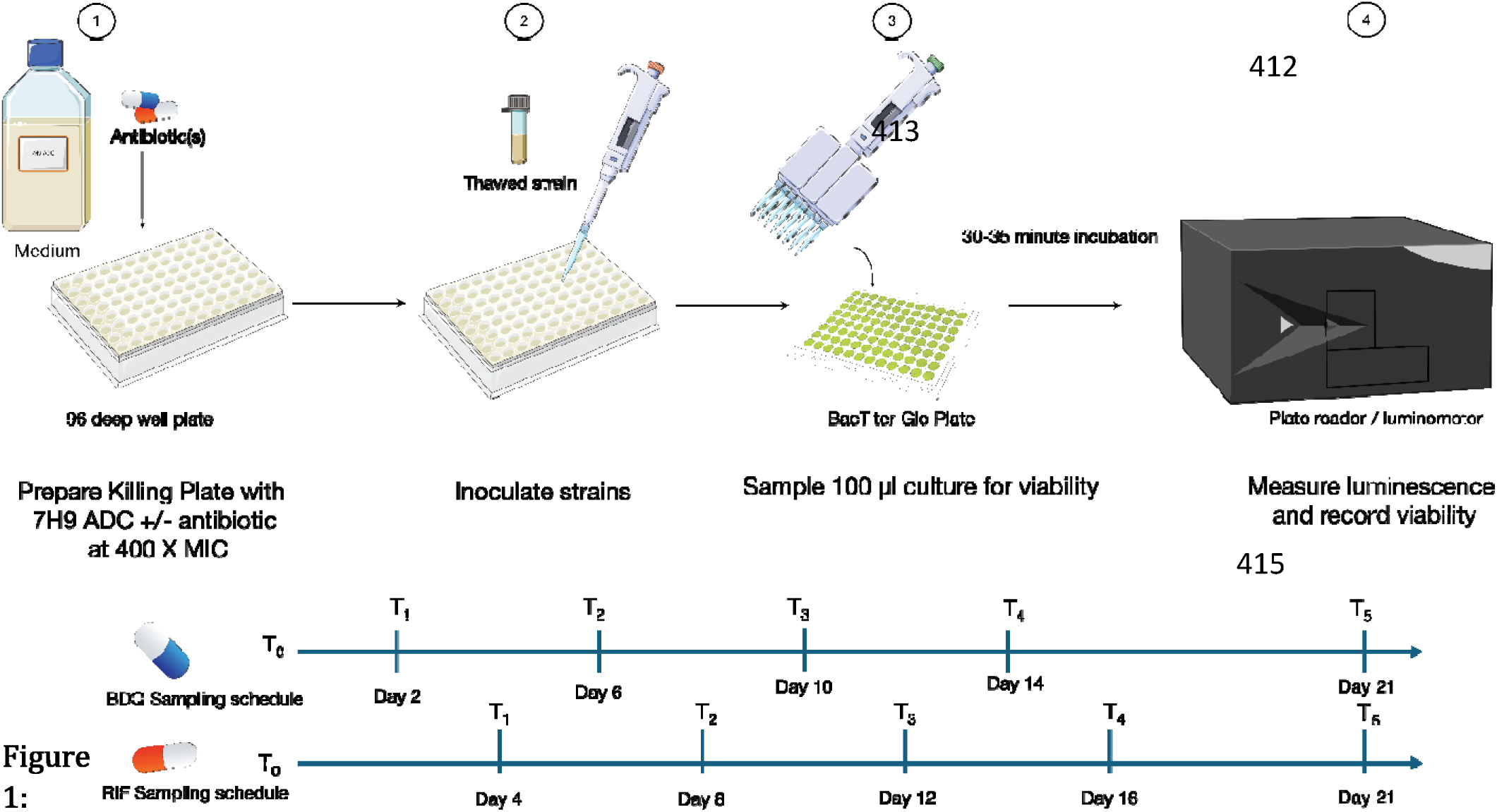
Experiment workflow for a real-time Time-Kill Assay (rt-TKA) to measure drug tolerance in Mtb (detailed in methods). The rt-TKA was set up following steps 1-4, and sampling occurred according to drug specific schedules following steps 3 and 4.

**Figure 2:**
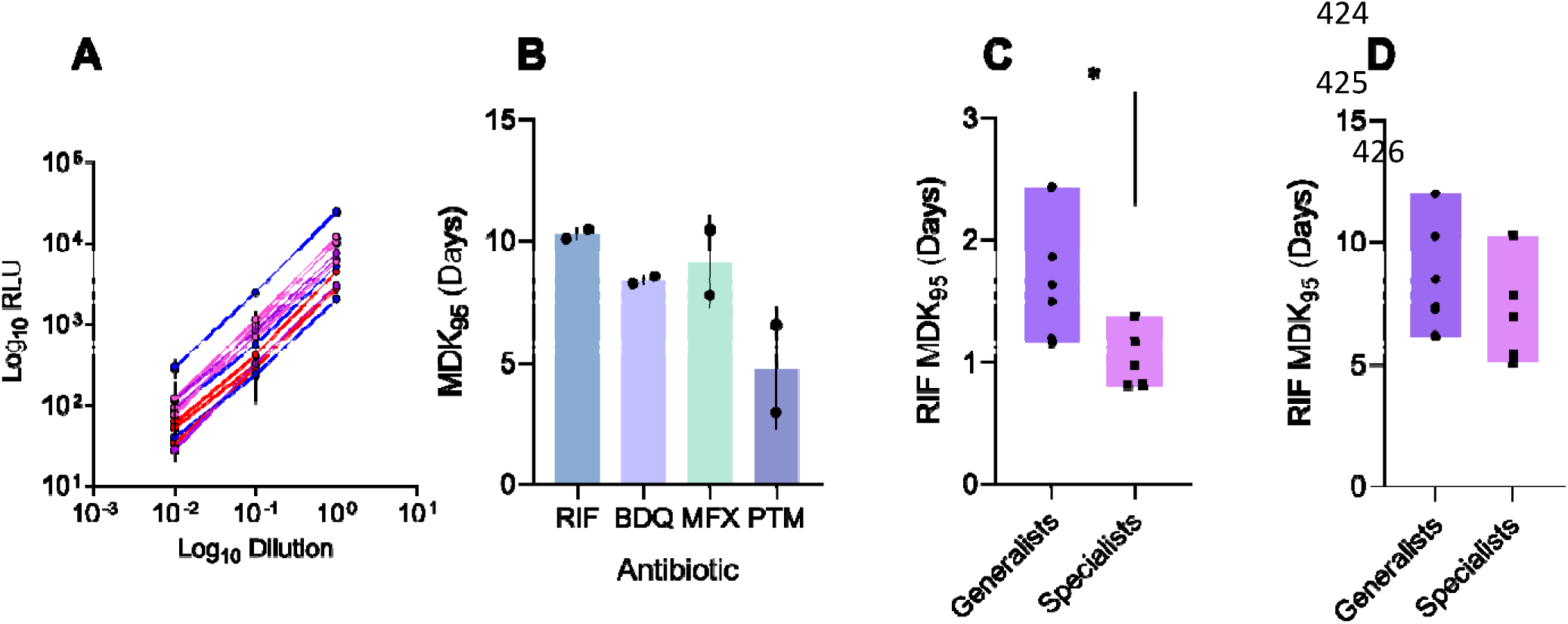
Benchmarking rt-TKA. **A.** Linearity of BacTiter-Glo ATP luminescence signal and bacterial density in reference strains of L1 – L4, n = 3 representative strains were inoculated into BacTiter- Glo in three log dilutions in triplicate. **B.** Reproducibility of minimum duration for killing (MDK_95_ values derived from rt-TKA in H37Rv exposed to rifampicin (RIF), bedaquiline (BDQ), moxifloxacin (Mfx) and pretomanid (PTM) at 100X MIC. Data derived from two independent experiments performed in triplicate. Bars denote mean, and error bars denote standard error. **C.** Comparison of RIF tolerance in reference generalist strains (L2 and L4, n = 6) and reference specialist strains (L1 and L3, n = 5) measured by CFU-based Time-Kill assay. Bars denote interquartile range. **D.** Comparison of RIF tolerance in reference generalist s (n = 6) and specialist strains (n = 5) measured by rt-TKA. Bars denote interquartile range. * p < 0.05 by t-test.

To check if the rt-TKA can recapitulate information from a standard approach, we compared tolerance in strains from geographically restricted lineages (L1 and L3), compared to globally distributed lineages (L2 and L4); or specialists and generalists, respectively. We compared MDK*_95_* values derived from a gold-standard CFU-based RIF time-killing assay at 25X MIC, to the RIF rt-TKA at 400X MIC (**Figure 2C and D**). We found that the rt-TKA could reproduce global patterns, but with less sensitivity. When comparing lineages, separately there were some deviations likely due to differences in experimental conditions such as drug doses and agitation (**Figure S1**).

### MIC distribution across a panel of Mtb clinical strains

We selected a genetically diverse panel of 102 drug-susceptible Mtb clinical strains spanning the globally most frequent lineages (L1 – L4). To confirm selected strains were susceptible to our drugs of choice, we measured strain minimum inhibitory concentrations (MICs) to the key front-line drug RIF and the key second-line drug BDQ. RIF susceptibility ranged between 0.06 µg/ml and 0.8 µg/ml, with a median of 0.05 µg/ml. BDQ susceptibility ranged between 0.31µg/ml and 1 µg/ml, with a median of 0.5 µg/ml.

Of the 102 strains assessed, two strains were found to be RIF-resistant based on the WHO critical concentration of 1 mg/L (**Figure 3A**). When comparing lineages, there was no specific pattern in RIF MICs (**Figure 3B**). For BDQ, three strains were found to be resistant which we defined as an MIC above but not including 1 µg/ml (**Figure 3C**). Here, strains belonging to L1 had significantly lower MICs than L3 and L4 (**Figure 3D**, p = 0.0088 and p = 0.0116, respectively), with members of sublineage L1.1.1 being significantly more susceptible (**Figure S3**). L2 strains also had lower MICs compared to L4 (p = 0.0295).

**Figure 3:**
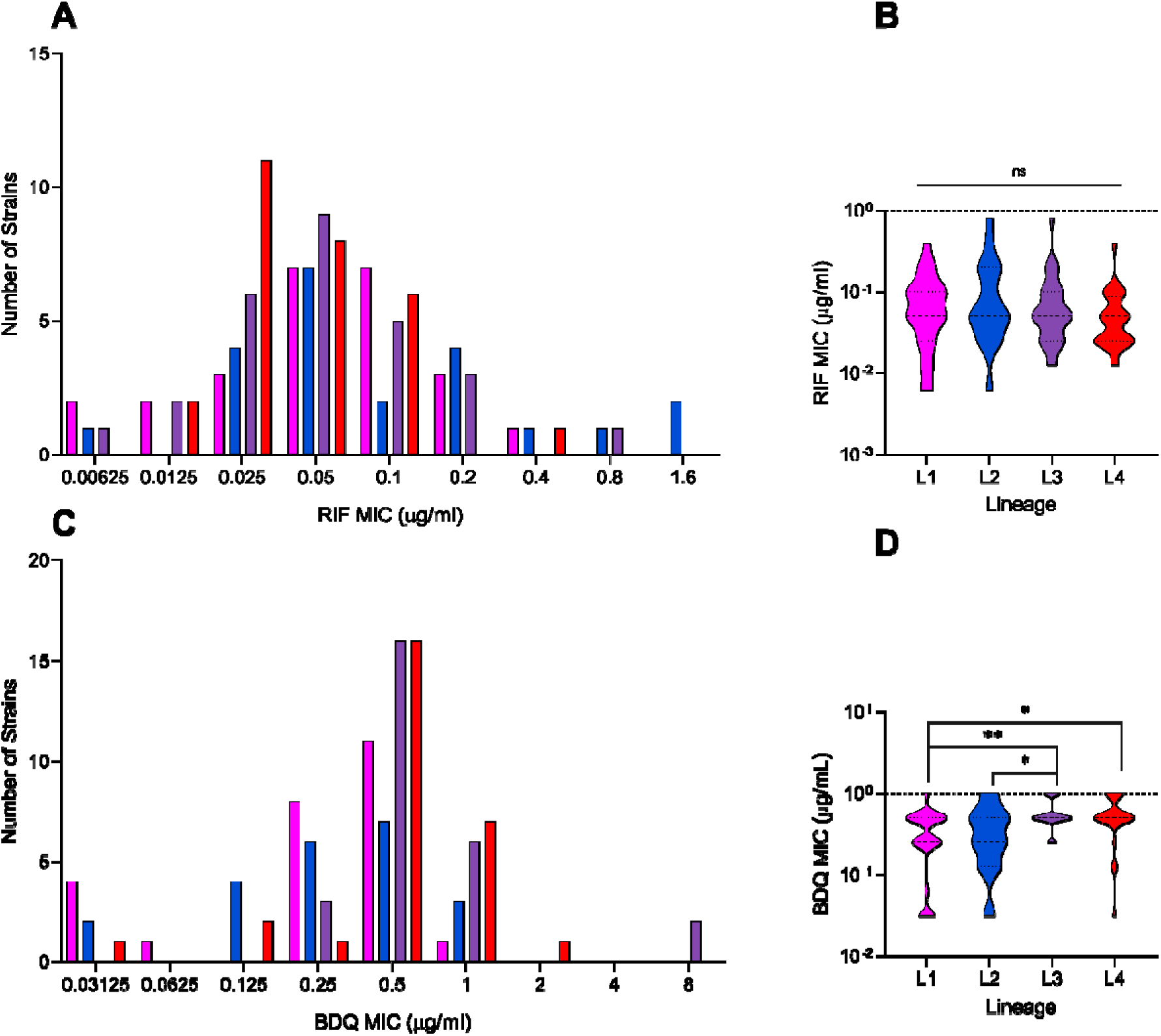
Drug susceptibility of a phylogenetically diverse strain panel representative of majo global Mtb lineages L1 to L4. **A**. Distribution of RIF minimum inhibitory concentration (MIC) in selected strains n = 102. Data derived from n = 2 independent measurements. **B**. Violin plots of MIC distributions in each lineage excluding resistant strains (n = 100). **C**. Distribution of BDQ MIC in strain panel, data derived from n = 2 independent measurements. **D**. Violin plots comparing tolerance by lineage in susceptible strains, * p < 0.05, ** p < 0.01 by Kruskal-Wallis with Dunn’s correction for multiple comparisons.

### Intrinsic drug tolerance in diverse Mtb strains varies across lineages

Using the rt-TKA, we measured tolerance to RIF and BDQ in the 99 drug-susceptible strains by deriving the minimum duration for killing 95% of the population (MDK_95_). We found tolerance to RIF ranged from 2.1 days to 15.5 days (**Figure S4A**), and tolerance to BDQ ranged from ∼1.2 to 14 days across strains (**Figure S4B**). In examining the differences in data distribution between the RIF and BDQ experiment, we noted a higher overall tolerance to RIF treatment compared to BDQ (**Figure 4**, p < 0.0001).

**Figure 4:**
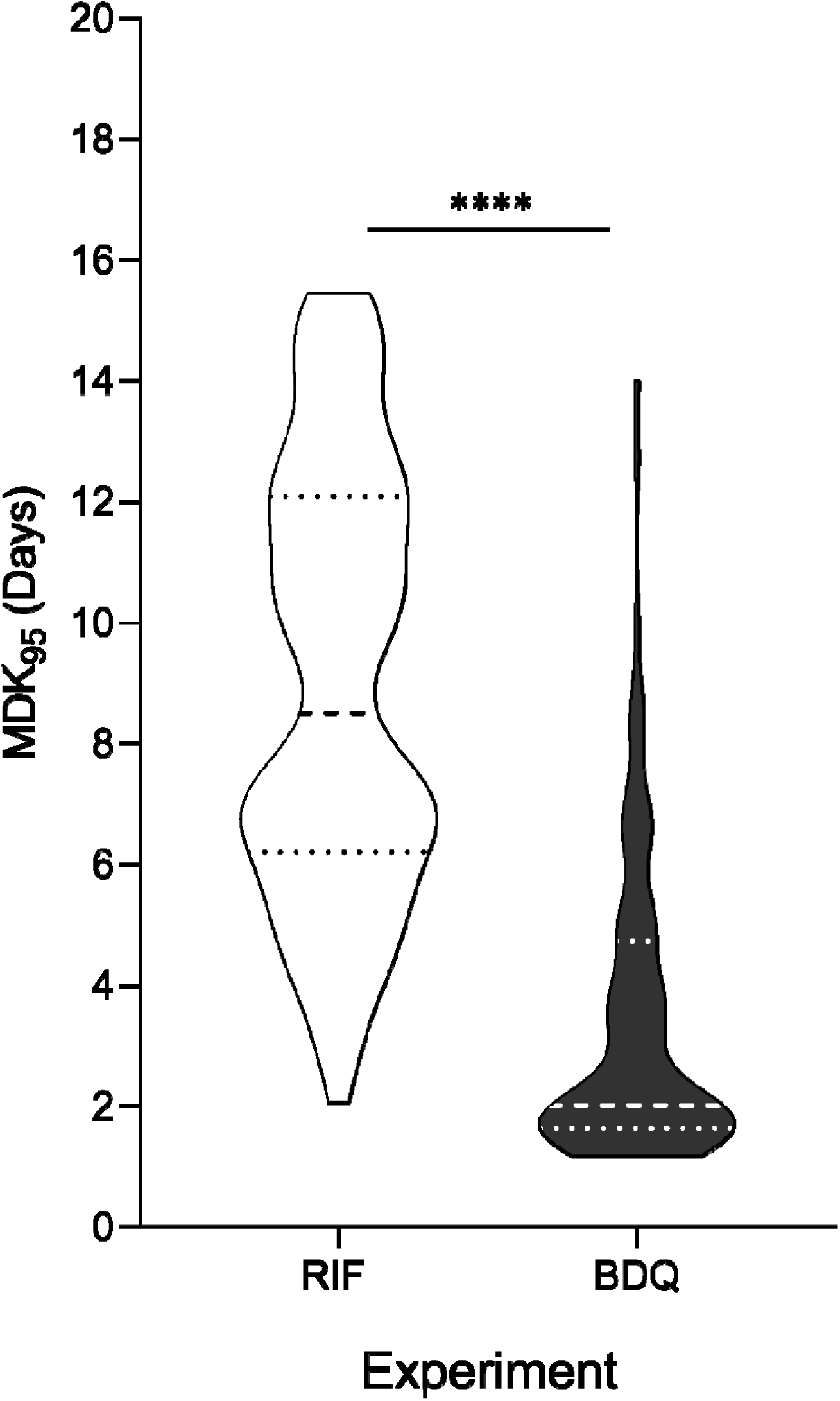
Violin plots comparing distributions of strain MDK_95_ values (n = 95) between RIF and BDQ experiments, dashed lines represent quartiles and median, **** p < 0.0001 by Kolmogorov- Smirnov.

To investigate the role of phylogenetic lineage in intrinsic tolerance to RIF and BDQ, we compared MDK_95_ values by lineage. When comparing tolerance to RIF by lineage, we found members of L1 to have significantly lower MDK_95_ values compared to L3 and L4 strains (**Figure 5A**, p = 0.0016 and p = 0.016, respectively). In addition, L2 strains had significantly lower MDK_95_ values and thus tolerance, compared to L3 strains (**Figure 3**C, p = 0.0313). We found stronger lineage associations when examining tolerance to BDQ, with L1 and L2 strains being significantly less tolerant compared to L3 and L4 (**Figure** 5B). We also found RIF tolerance to have an inverse relationship with MIC, whereas the opposite was true for BDQ, where strains with higher MICs had a higher tolerance in general (**Figure S8**). Thirteen strains had high tolerance (upper quartile in both instances) to both BDQ and RIF (**Figure S**7), indicating potential multidrug tolerance.

**Figure 5:**
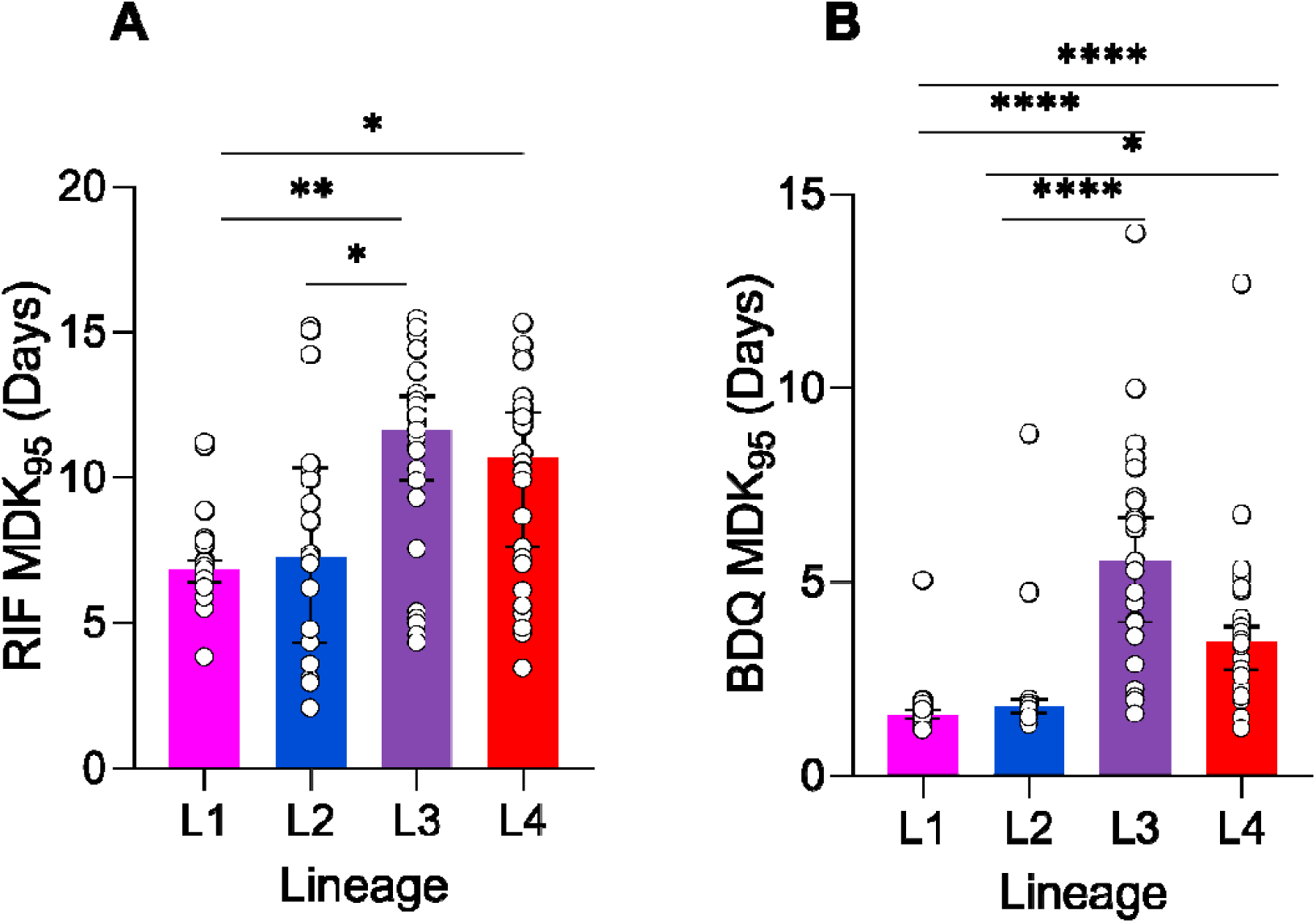
Lineage specificity for high tolerance to RIF and BDQ. **A.** Comparison of RIF tolerance between L1 (n = 25), L2 ( n = 19) L3 ( n = 27) and L4 ( n = 28) strains. Bars denote median, with 95% confidence interval (C.I.) error bars. * p < 0.05, ** p < 0.01, **** p < 0.0001 by Kruskal-Wallis with Dunn’s correction. **B.** Comparison of BDQ tolerance between L1 (n = 25), L2 (n = 22) L3 (n = 25) and L4 (n = 26) strains. Bars denote median, with 95% C.I.

### Tolerance-associated genetic markers in diverse strain set

Having generated a phenotypic dataset of tolerance to two drugs, we ventured to explore genotypic associations that could account for our phenotypic data. First, we wanted to assess the heritability of MIC and MDK_95_ in our strains, andunderstand if genetically similar strains would exhibit a similar level of tolerance. To that end, we constructed a maximum likelihood tree of our strain set and plotted levels of tolerance at the relevant tips. There were clearly similar levels of tolerance in phylogenetically more closely related strains by visual inspection (**Figure 6**). We further computed phylogenetic signals in our data using Pagel’s Lambda and Blomberg’s K and found significant signals for MIC and MDK_95_ (**Table 1**). From positive control (drug-free) wells, we were able to extrapolate relative growth in our strains (**Supplementary methods, Figure S9**), which, in contrast to MIC and MDK_95_ values, did not show significant heritability (**Table 1**).

**Figure 6:**
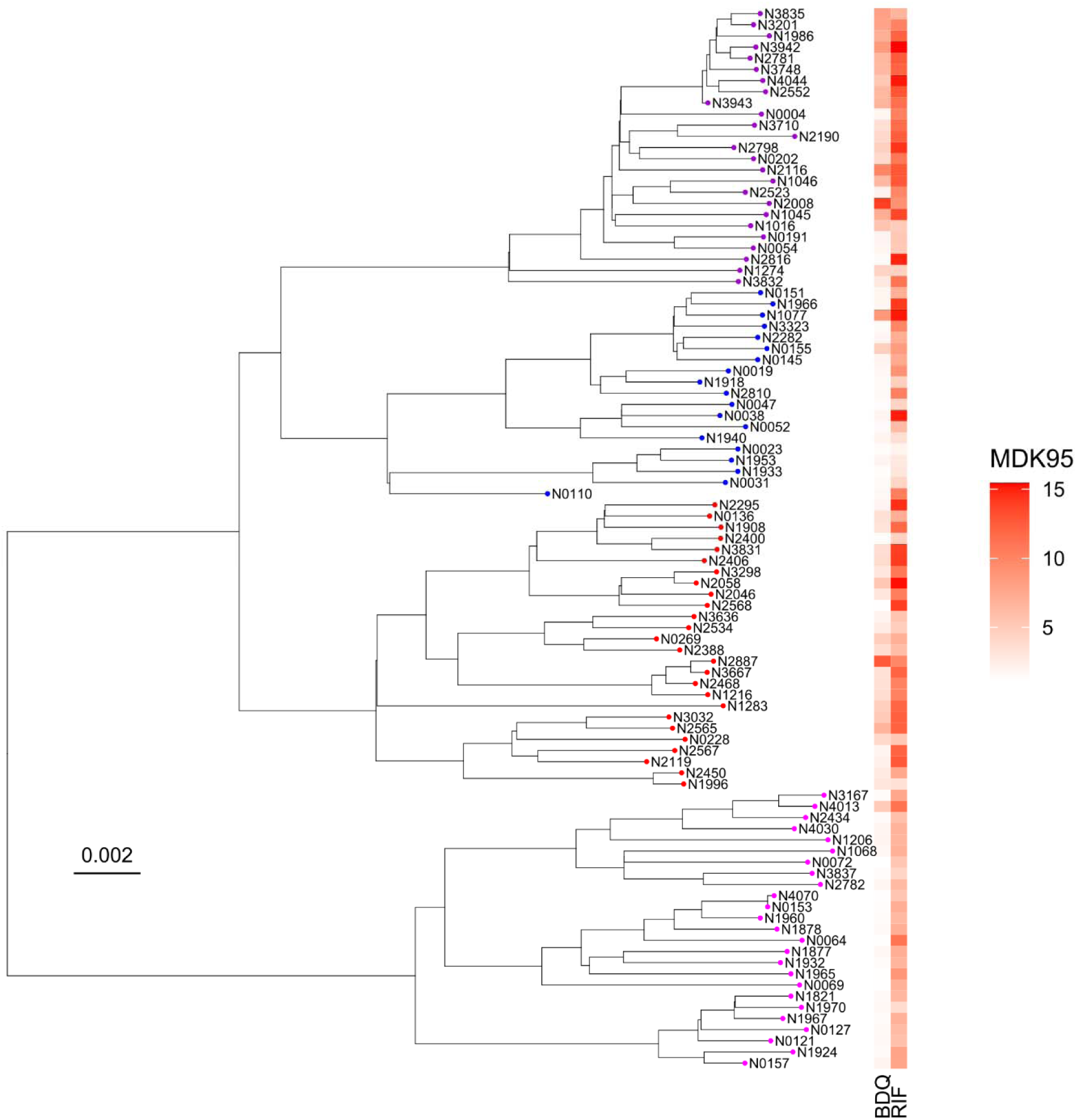
Maximum likelihood phylogeny-based representation of the Mtbstrain set and corresponding level of tolerance to RIF and BDQ. Tips coloured by Mtb lineage L1 = pink L2 = blue L3= purple L4= red. Colour scale depicts level of tolerance according to rt-TKA data. Scale bar represents substitutions per polymorphic site.

**Table 1:**
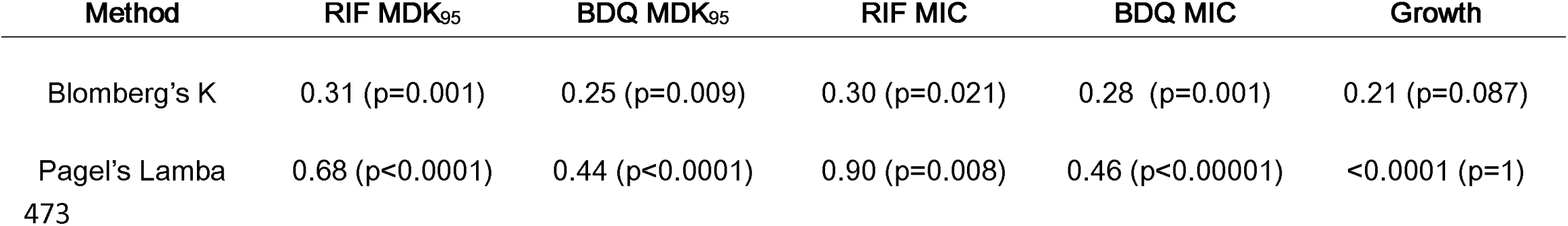
Heritability analysis using phylogenetic signalling of MDK_95_, MIC, and Growth in strain set.

Next, we attempted to uncover specific mutations that could account for our data; we performed a genome wide association study (GWAS) with tolerance as a continuous variable for both sets of data. Overall, we found no significant polymorphisms based on the set threshold of 2.46 e-5 (see methods). However, there were polymorphisms common to strains exhibiting high tolerance to BDQ in a set of L3 strains. (**Figure S12-14, Table S1**). None of the identified positions corresponded with known genes associated with drug tolerance.

We compiled a list of genes that have previously been associated with drug tolerance in the literature (**Table S2**). To be as inclusive as possible, in addition to tolerance-associated genes, we considered loci related to drug persistence and resilience (Refs). Using our tolerance data, we performed an enrichment analysis for polymorphisms associated with level of tolerance to RIF and BDQ (**Table 2**). We designated upper quartile strains for tolerance to RIF and BDQ as high tolerance strains and performed an enrichment analysis for both experiments (Methods). The P271T mutation in PflA (Rv3138) was significantly enriched in RIF tolerant strains. Many mutations were enriched for in high BDQ tolerant strains, which mainly are associated with lipid metabolism, cell wall and cell process, intermediary metabolism and information pathways (**Table 2**). However, most identified mutations were private to L3 strains (**Figure S15**). By contrast, Tig (Rv2462c) L118P, LppD (Rv1899c) P123L, and FadD26 (Rv2930) V240M occurred multiple times in multiple lineages but were less frequent or absent in L1 strains (**Figure S15**).

**Table 2.**
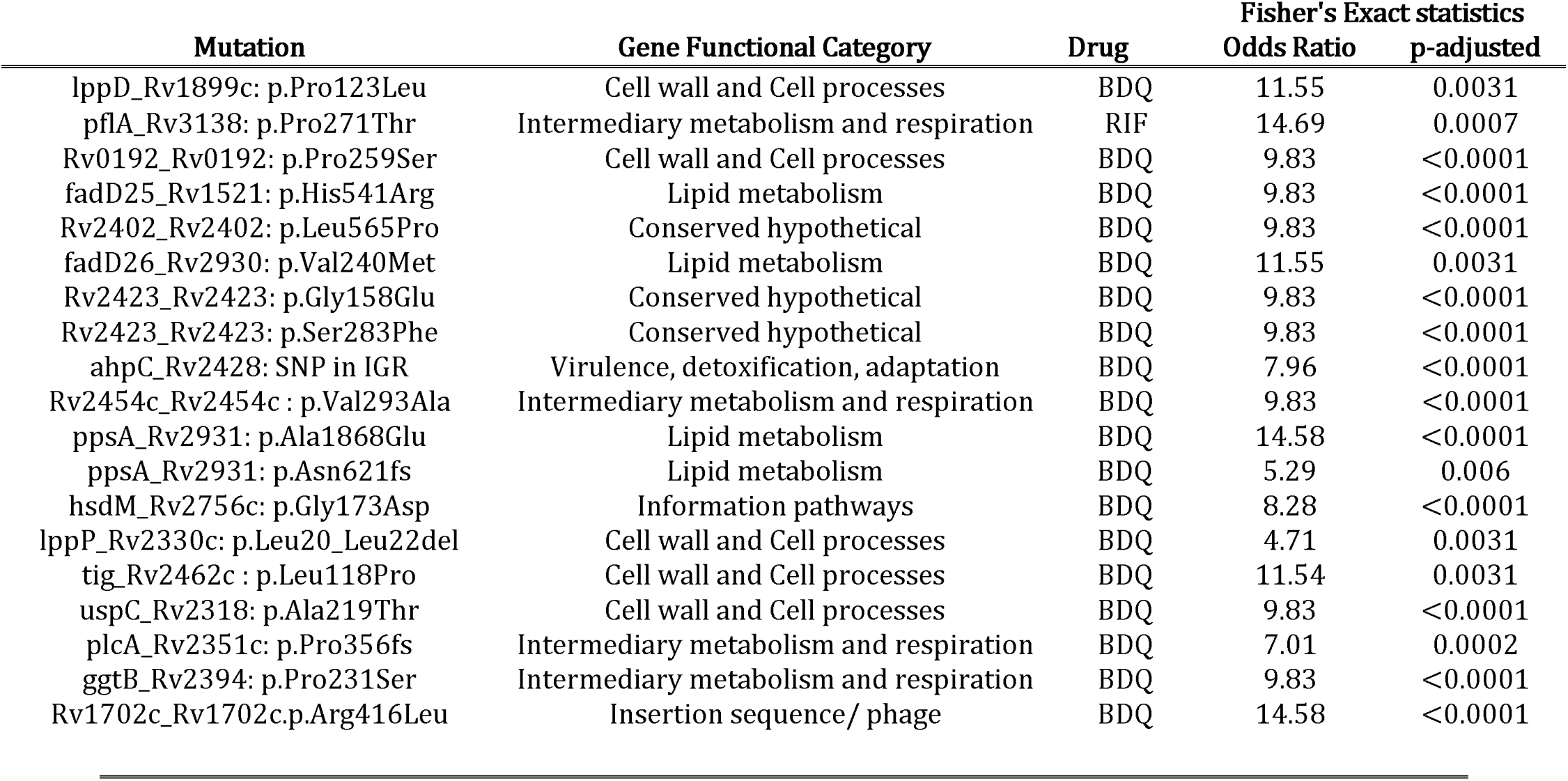
Mutations significantly associated with upper quartile tolerance in strain set.

## Discussion

MDR-TB outbreaks are frequently caused by strains belonging to L2 and L4 of the MTBC. We hypothesized that L2 and L4 have higher intrinsic drug tolerance and thus more readily evolve resistance. To investigate this, we devised an *in vitro* assay to measure drug tolerance in Mtb. The assay exposes bacteria to killing concentrations of antibiotics dosed based on strain-specific MICs. We selected a panel of phylogenetically diverse clinical strains to interrogate whether MTBC lineage relates to tolerance to RIF and BDQ. Drug susceptibility testing for RIF and BDQ revealed that L1 strains have higher BDQ susceptibility, while RIF susceptibility showed no lineage association. Strains were generally more tolerant to RIF compared to BDQ. By lineage, L1 strains had low tolerance, and L3 and L4 strains had higher tolerance overall to both drugs. Using phylogenetic signalling methods, we found genetically similar strains to have similar levels of tolerance, which was also true for strain MICs. Genetically, we found statistically significant enrichment for mutations in genes involved in stress response, virulence and cell wall architecture in high tolerance strains.

We adapted existing strategies using intracellular ATP for our time-kill assay, utilizing the Promega BacTiter-Glo assay to measure viability across time (32–36), which we dubbed the rt-TKA. In doing so, increasing throughput, with 100 strains measured in parallel, real-time and in a single experiment under BSL3 limitations. From the initial hypothesis, we expected to see higher tolerance in “generalists” (L2 and L4) compared to “specialists” (L1 and L3) which was true in both CFU and rt-TKA data. Thus, recapitulating similar findings in assays with different dosing (25 X MIC vs 400 X MIC) and oxygen availability (shaking falcon tubes vs static deep well plates) indicated that the rt-TKA could retrieve biologically consistent data. Moreover, regardless of lineage, our assay could detect decreases in density up to two log fold, appropriate for measuring time kill kinetics for tolerance according to convention (37, 38).

With higher throughput, we selected strains belonging to global majority MTBC lineages L1 – L4. Within each group, we selected strains from subclades that are globally prevalent (**Supplementary data**) (39–41) and also included more geographically restricted sublineages in L2 and L4, namely L2.1 and L4.6, respectively. In susceptibility testing, consistent with a previous report, L1 strains were significantly more susceptible to BDQ (42). Otherwise, the distributions for RIF and BDQ MICs corresponded with known susceptibility profiles in circulating strains (43–47).

Prior to our work, variation in RIF tolerance in Mtb had been established (48, 49). This tolerance could be a property of recent adaptation shaped by TB treatment in patients (48). Alternatively, it could be an intrinsic property based on the macroevolutionary events that define MTBC lineages, giving rise to differentially drug adaptable phenotypes. We found that overall, strains exhibited higher tolerance to RIF compared to BDQ, which may speak to a role for recent adaptation as BDQ is a newer drug. However, L3 and L4 strains were more tolerant to both RIF and BDQ compared to L1; two different drugs with two different modes of action, suggesting that features patterned by phylogeny rather than recent adaptation may also play a role. Some strains exhibited high tolerance to both RIF and BDQ, indicating the possibility of a multidrug-tolerant phenotype, alongside drug-specific tolerance. This calls for a deeper exploration of the specific phenotypic attributes that govern drug tolerance in these different circumstances.

Probing into a molecular basis for drug tolerance in our dataset, we explored genetic features. With phylogenetic signalling, we found tolerance to be a heritable trait (50), corresponding with a recent study lin the nontuberculous mycobacterium *M. abscessus* (51). Using our tolerance data for a GWAS was unsuccessful, likely due to sample size. However, we noted polymorphisms that could be of interest. We further investigated genes previously associated with tolerance or the related phenotypes ofdrugpersistence and resilience (52), to see if there were specific mutations enriched in high-tolerance strains.

High RIF tolerant strains had significant enrichment for the P271T mutation in PflA (Rv3138). PflA is has previously been shown to be upregulated in stationary phase (53) and has been associated with RIF tolerance (54). For BDQ that had more associated polymorphisms, including mutations in *ppsA*, which is involved in phthiocerol dimycocerosate (PDIM) synthesis, a virulence lipid associated with persistence and drug response, including drug tolerance (55–57). There was also enrichment for lipoproteins (*lppP, lppD*) and lipid metabolism (*fadD25*, *fadD26*, *plcA*), alongside several genes associated with stress response (*ggtB*, *ahpC*), and metabolism (Rv2454) (54). Most of these mutations occurred exclusively in L3 strains, suggesting that L3 adaptation has resulted in a phenotype especially tolerant to BDQ, however more evidence is required for this assertion. Outside of L3 alone, mutations in protein chaperone Tig, lipoprotein LppD, and fatty acyl-AMP ligase FadD26 were associated with BDQ tolerance. Collectively, our results suggest that adaptation in systems involved in oxidative stress, metabolism and cell wall architecture could impart drug tolerance in Mtb.

Our study has some limitations. While the method developed here, we did not include a washing step during sampling, which allows any free-floating extracellular ATP to be included in measurements. In theory, the proportion of this extracellular ATP should be based on the proportion of bacteria that died which should not bias the outcome, but an apyrase treatment could be implemented to overcome this (58). There could be a risk that the lysis agent in the BacTiter-Glo reagent did not fully lyse the entire sample. However, Ferriera et al. showed that additional mechanical disruption did not yield more ATP than BacTiter Glo alone in Mtb (34). In our study, we saw that a peak luminescence signal was reached after 20 minutes of incubation on average with no specific difference in lysis in the BacTiter-Glo reagent based on lineage (**Figure S1**). Based on our benchmarking, the assay was less sensitive than the CFU standard, but our assay reproduced previous results by others (Refs). This includes RIF tolerance being negatively associated with RIF MIC (54), an association between PflA polymorphisms and RIF tolerance (48), and higher tolerance in low ATP producers (59–61). Additionally, the rt-TKA does not rely on regrowth to measure viability, hence being “real time”. Assays such as the colony forming unit assay or most probable number assay require regrowth in solid or liquid media respectively for quantification of viable bacteria. This regrowth has been shown to affected by the bacterial drug response, wherein the capacity to regrow in media can be affected, thus jeopardizing the interpretation of subsequent killing data (22, 62), which our assay avoids.

In conclusion, we have devised a convenient, high throughput method to study drug tolerance in Mtb, and interrogated whether strains of L2 and L4 are more intrinsically drug tolerant, given their repeated association with drug resistance (15, 63). We found L2 strains to not have significantly high tolerance in our experimental setting while L4 strains exhibited high tolerance alongside L3. This could suggest that drug tolerance might contribute to resistance evolution in a lineage-dependent manner. L3 strains are not typically associated with MDR-TB outbreaks, but drug-resistant L3 strains do emerge (64). However, they are not as globally successful as their L2 and L4 MDR counterparts(10). Taken together, our data show that there is variability of tolerance to two key anti-TB drugs across Mtb strains and that this variability has a lineage association. What remains to be understood is the epidemiological relevance thereof, for which we have established a foundation from which to further explore.

## Acknowledgements

This work was supported by the Swiss National Science Foundation (grants 320030-227432, 320030L-231163 and CRSII5_213514) and the European Research Council (883582- ECOEVODRTB). Calculations were performed at sciCORE (http://scicore.unibas.ch/) scientific computing core facility at the University of Basel. Sequencing was carried out at the Genomics Facility Basel of the University of Basel and the Department of Biosystems Science and Engineering at ETHZ in Basel, Switzerland. Credit to servier for vector graphics https://smart.servier.com/ licensed under CC-BY 3.0 Unported https://creativecommons.org/licenses/by/3.0/.

## Author Contributions

Study conception: V.F.A.M., D.B., S.Bor., and S.G ; experimental design: V.F.A.M., D.B., C.L., S.Bor., and S.G.; data acquisition and analysis: V.F.A.M., M.Z., D.B., C.L., G.A.G.,S.Bou., S.K., A.D., and M.R.; interpretation of data: V.F.A.M., D.B., S.Bor., and S.G.; drafted the main manuscript text: V.F.A.M., S.Bor., and S.G.

## Code and Data Availability

All data are included within the main text and supplementary materials. Code available at: https://git.scicore.unibas.ch/TBRU/mdkcalculator. Genomic sequences were uploaded under bioproject: PRJEB77138.

