## Supplementary Information for "Variability in intrinsic drug tolerance in *Mycobacterium tuberculosis* corresponds with phylogenetic lineage"

**Supplementary Methods:**

**BacTiter-Glo incubation Test:**

Reference Mtb strains were inoculated into 7H9 ADC in triplicate in 96 deep well plates according to the rt-TKA protocol. After gently mixing, 100 ul was sampled and inoculated into BacTiter-Glo plates, which were sealed with plastic film and disinfected for removal from BSC. After to 5 minute incubation, plates were immediately placed into Tecan Infinite Pro for luminescence measurement. The plate reader was programmed to take a measurement every 5 minutes for 120 minutes.

**Extrapolating relative growth**

Fold change relative light unit (fcRLU) values were calculated from positive control wells of rt-TKA experiment. Values for each strain were plotted to identify time points where linear exponential growth occurred. The slope of resulting growth curves was calculated by dividing the change in fcRLU in the interval by the change in time in the interval to derive a relative growth metric. An average relative growth was determined by combining calculations from RIF and BDQ experiments.

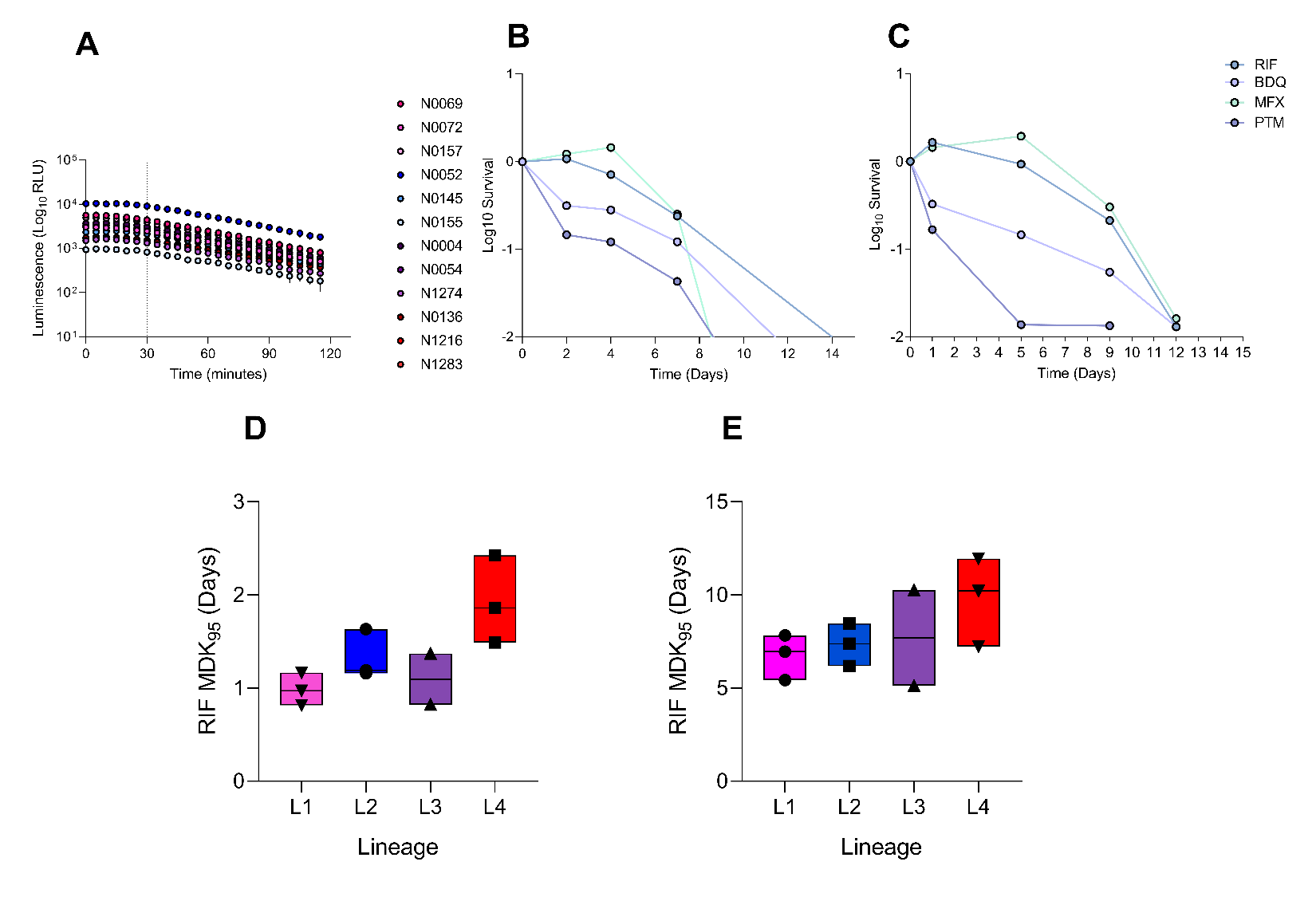

**Figure S1**: rt-TKA optimization. **A.** Change in luminescent signal from BacTiter-Glo incubation across time in three representative strains from L1 – L4. **B.** H37Rv rt-TKA pilot scheduled for sampling on day 2, 4, 7 and 14. **C.** H37Rv rt-TKA pilot with sampling scheduled on day 1, 5, 9 and 12. **D.** Classic tolerance assay sampled by CFU, separated by lineage. **E.** rt-TKA of same strains, separated by lineage.

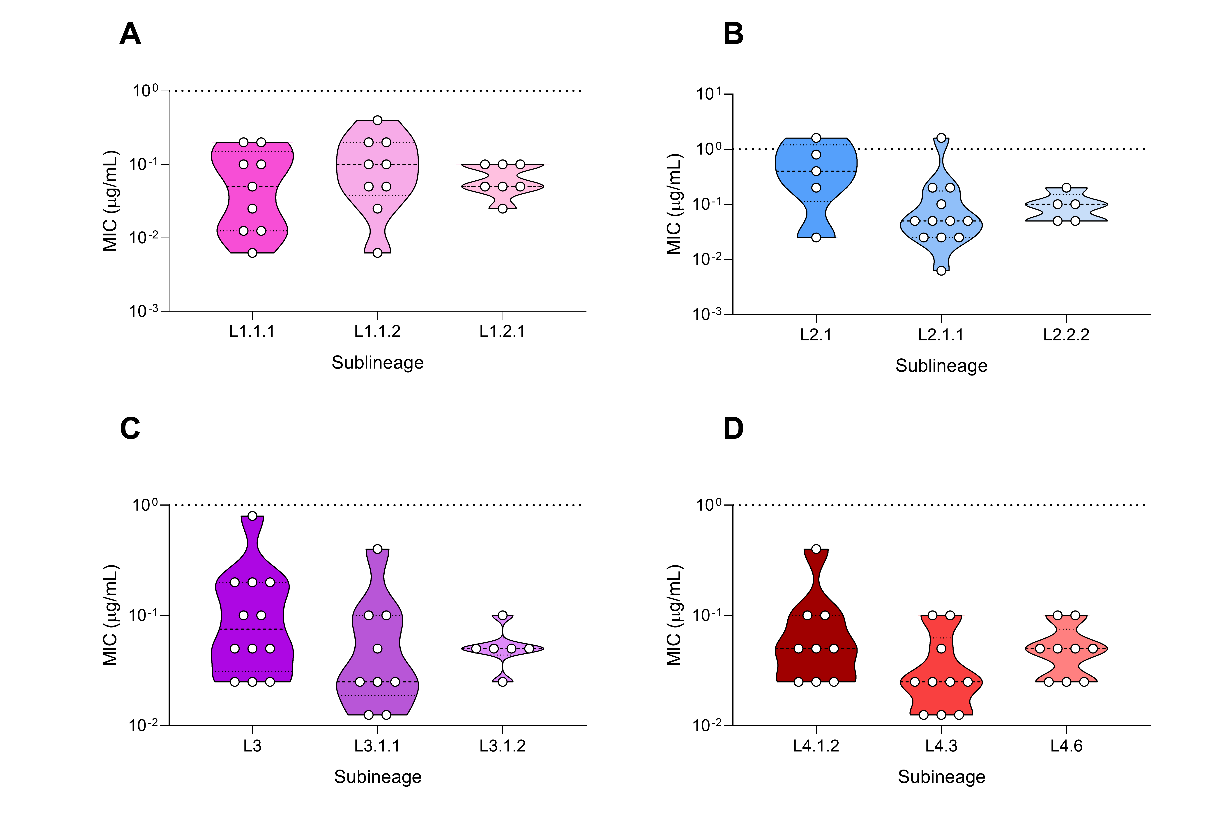

**Figure S2:** Violin plots of RIF MICs by sublineage. **A.** MICs in L1 strains. **B.** MICs in L2 strains. **C.** MICs in L3 strains. **D.** MICs in L4 strains.

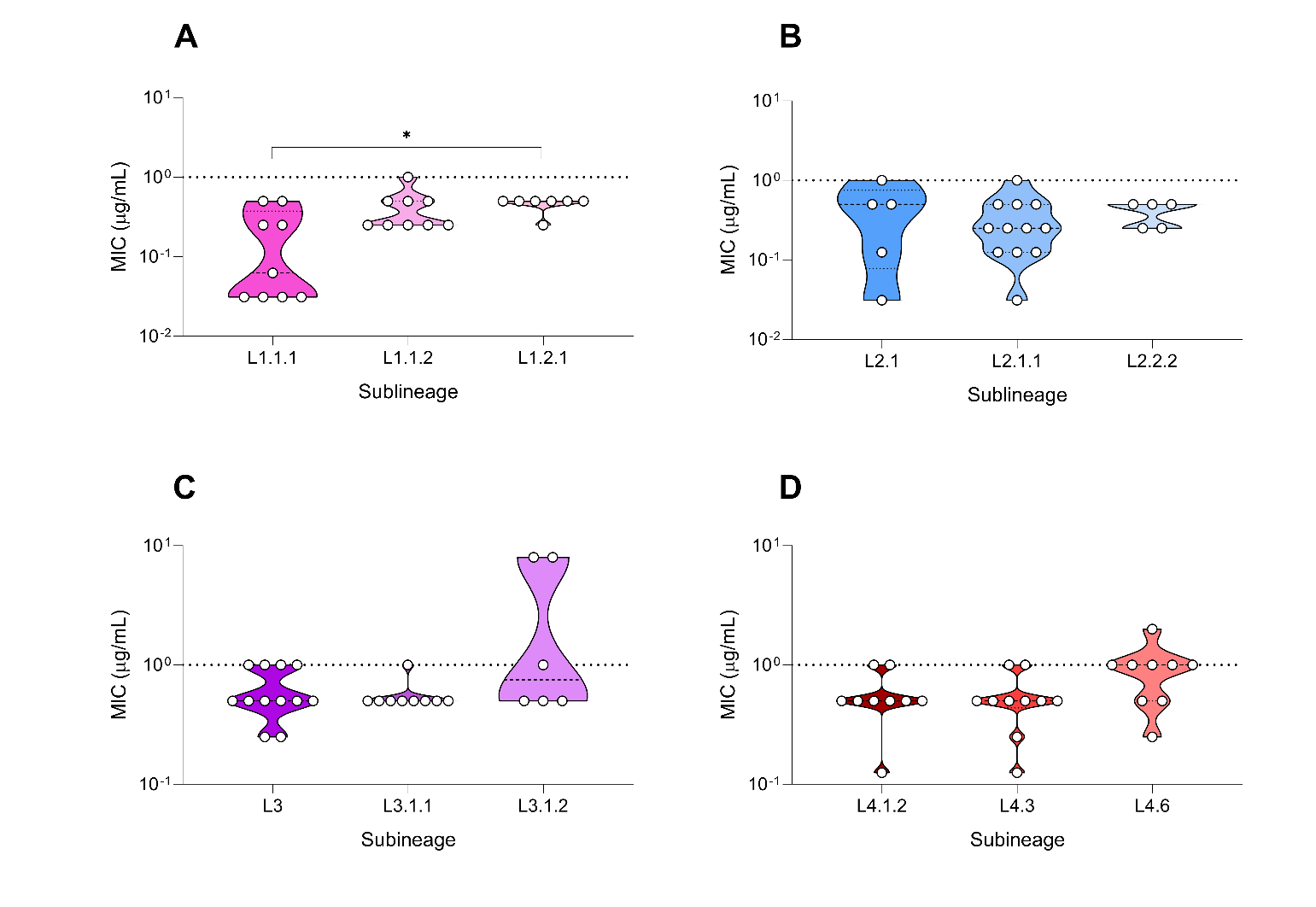

**Figure S3:** Violin plots of BDQ MICs by sublineage. **A.** MICs in L1 sublineage strains. **B.** MICs in L2 sublineages. **C.** MICs in L3 sublineages. **D.** MICs in L4 sublineages. * p = 0.0167 by Kruskal-Wallis with Dunn’s correction for multiple comparisons.

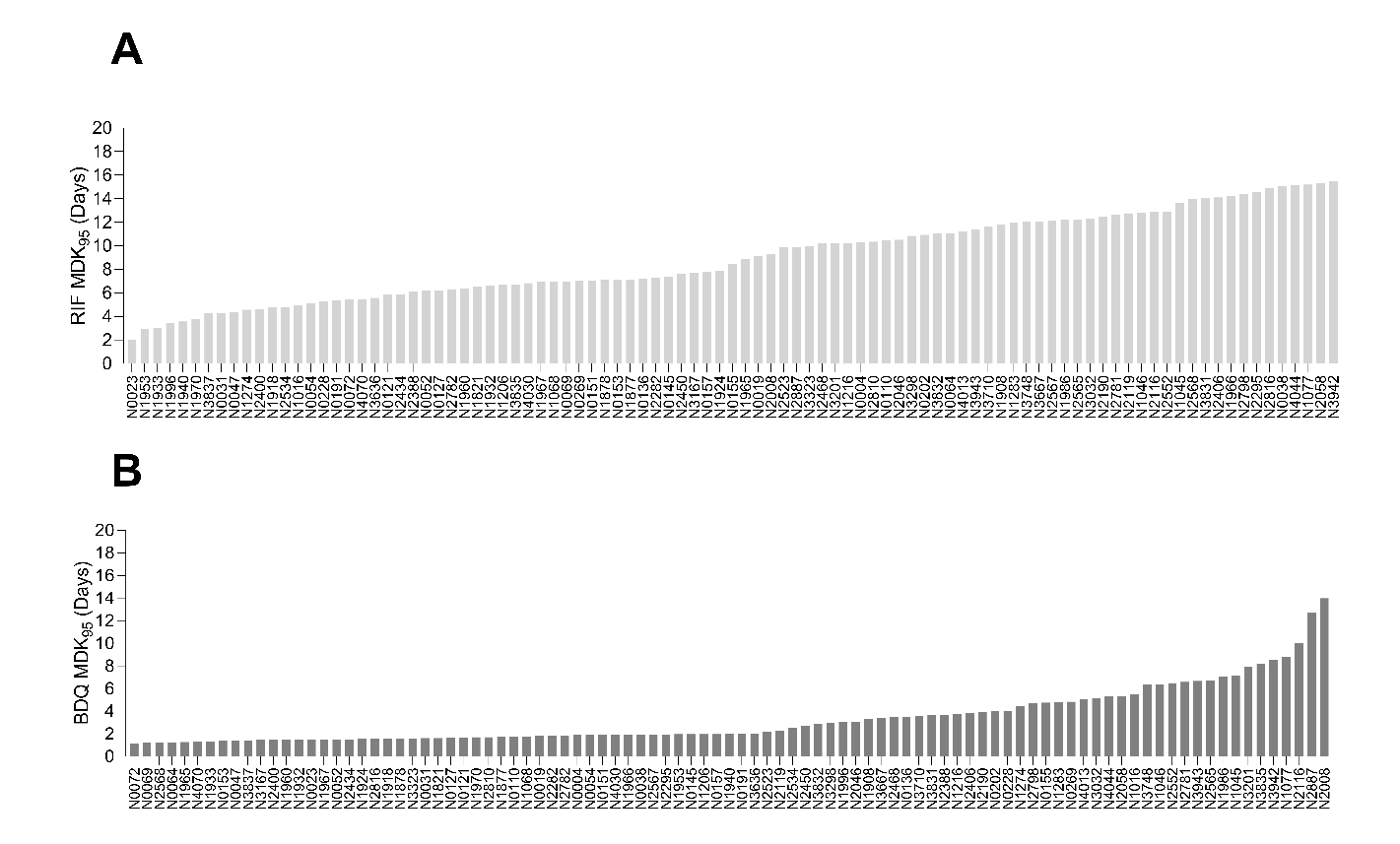
**Figure S4:** Tolerance in diverse strains of Mtb (n = 95). **A.** Distribution of RIF tolerance (n = 95) in strain set exposed to 400 X MIC RIF, measured by rt-TKA. X-axis indicates strains. Data representative of one independent experiment performed in triplicate. **B.** Distribution of BDQ tolerance (n = 95) at 400 X MIC, measured by rt-TKA. Data representative of one independent experiment performed in triplicate

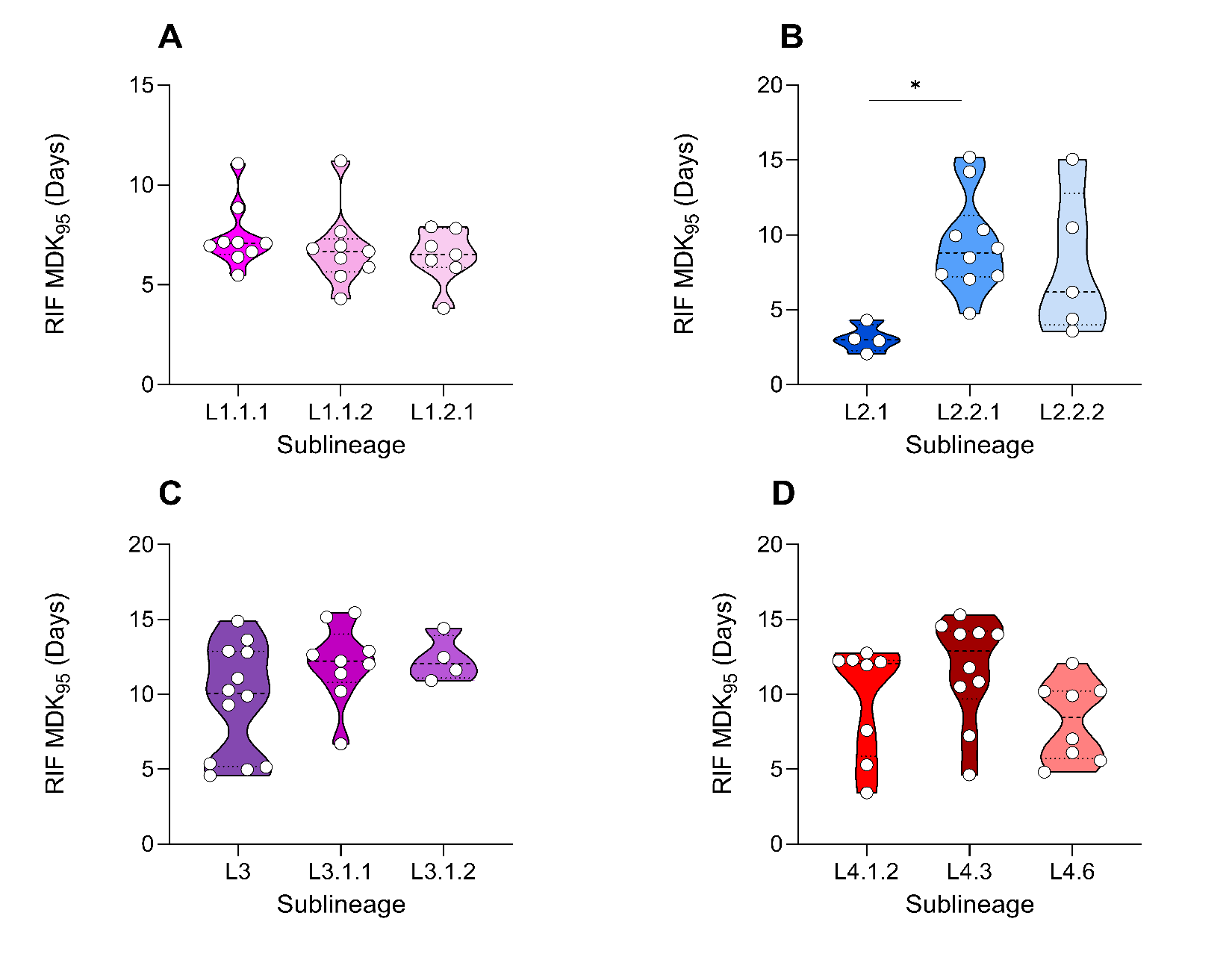

**Figure S5:** Violin plots displaying tolerance to RIF by sublineage: **A.** Tolerance in L1 strains. **B.** Tolerance in L2 strains. **C.** Tolerance in L3 strains. **D.** Tolerance in L4 strains. * p = 0.0182 by Kruskal-Wallis with Dunn’s correction for multiple comparisons.

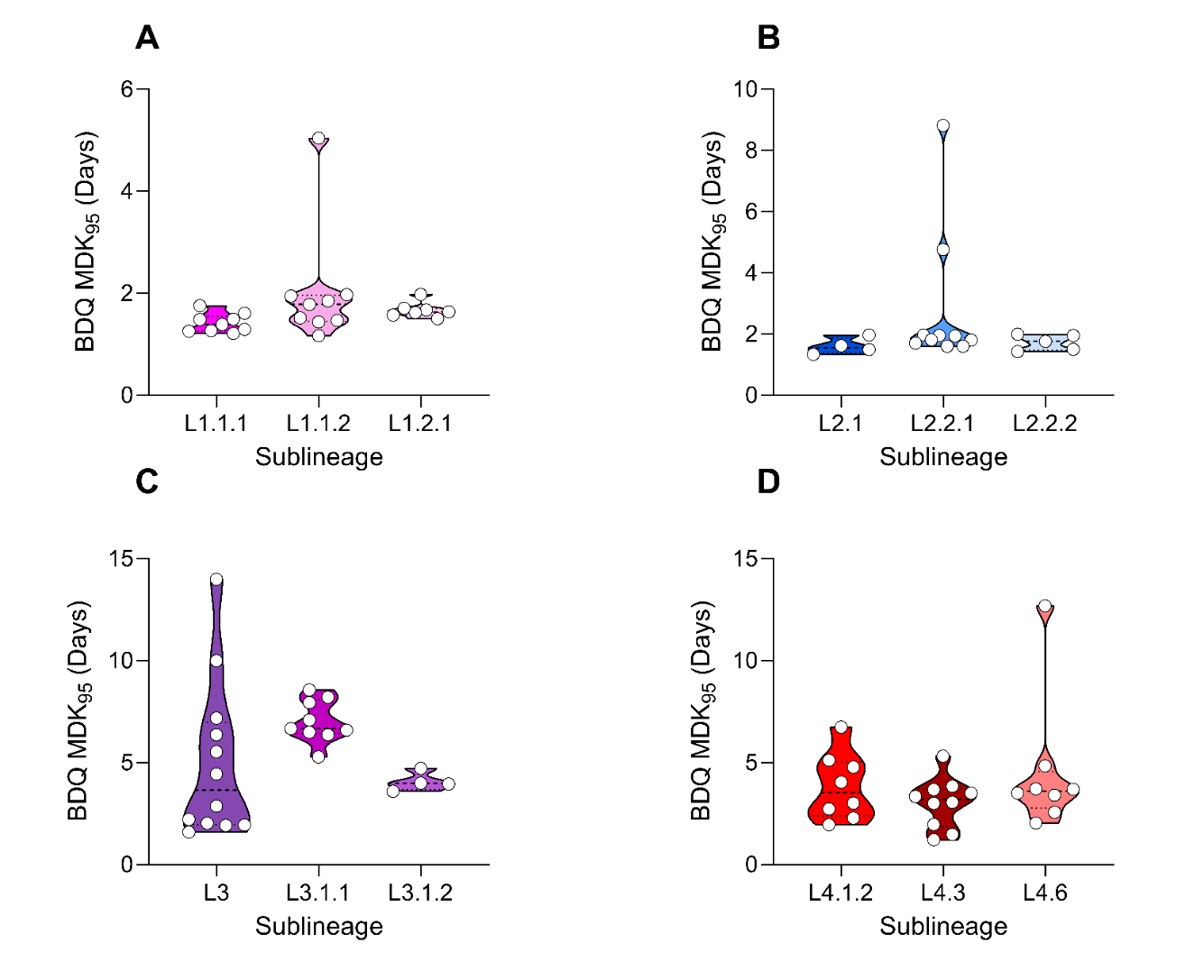

**Figure S6:** Violin plots displaying tolerance to BDQ by sublineage: **A.** Tolerance in L1 strains. **B.** Tolerance in L2 strains. **C.** Tolerance in L3 strains. **D.** Tolerance in L4 strains.

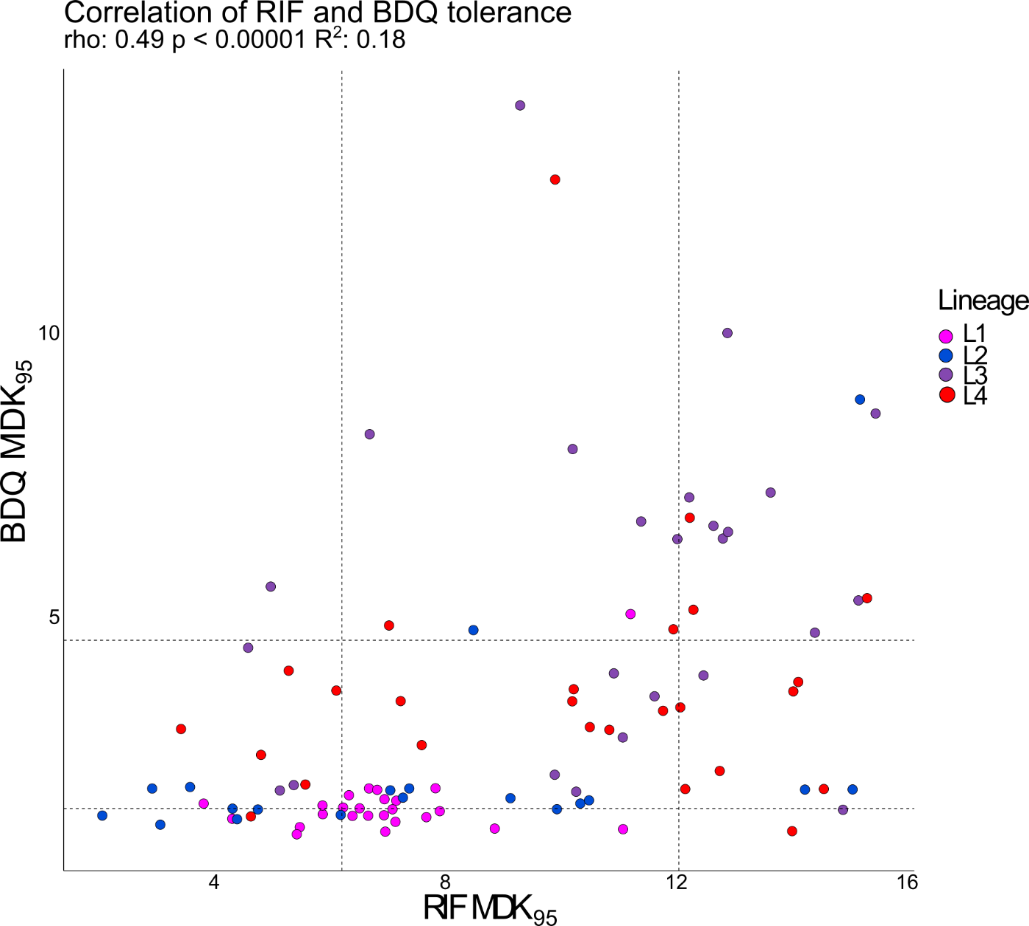

**Figure S7:** Projection of tolerance in strains in RIF and BDQ experiments. Dashed lines indicate upper and lower quartile values for MDK_95_s in for RIF (vertical lines) and BDQ (horizontal lines).

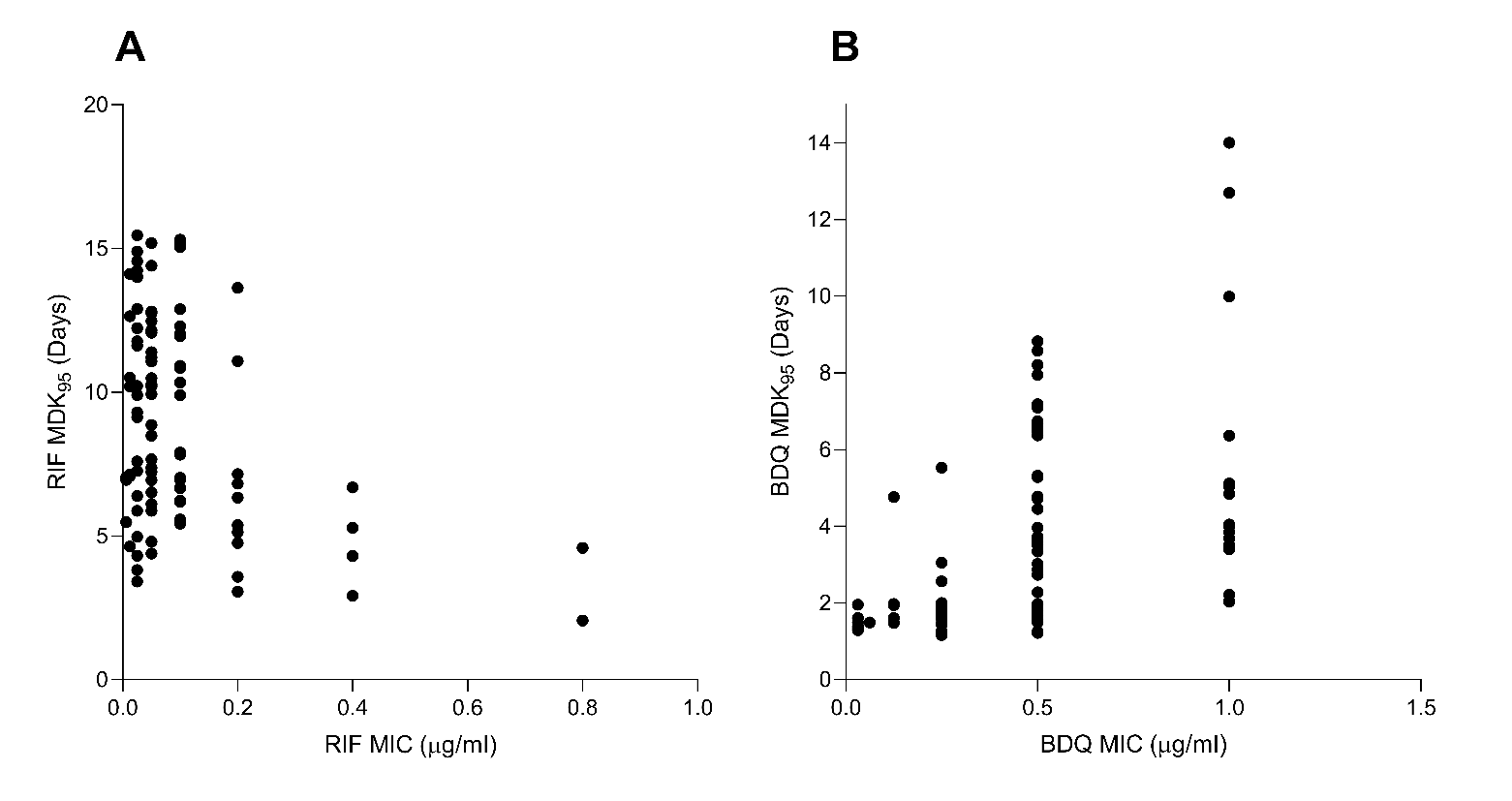

**Figure S8:** Projection of MIC and MDK95 for each drug. **A.** Inverse relationship between RIF MIC and RIF MDK_95_. **B.** Positive relationship between BDQ MIC and BDQ MDK_95_.

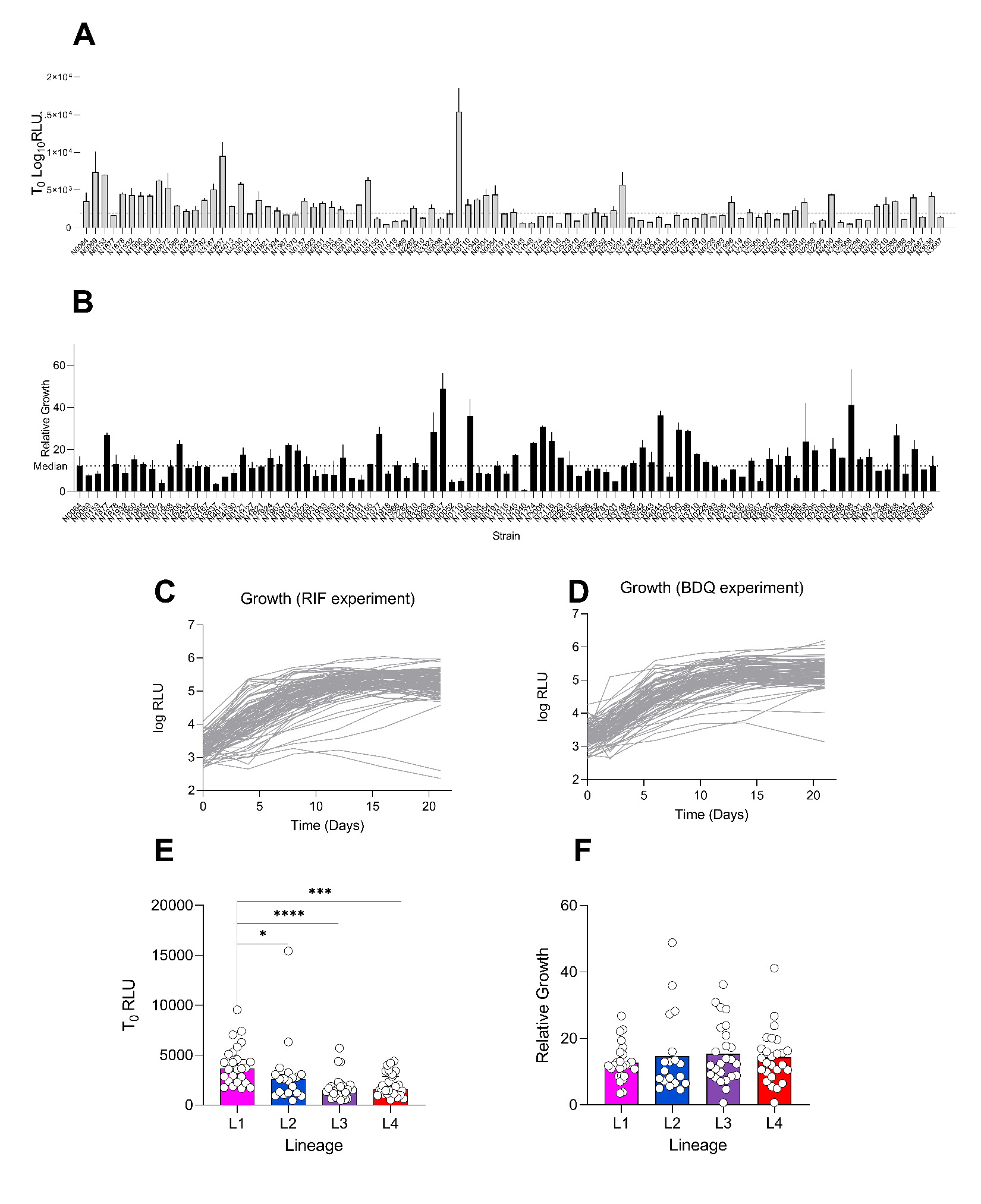

**Figure S9:** Non-drug-related metrics extracted from rt-TKA data. **A.** Initial ATP signal. Bars denote mean initial measurement in each strain extrapolated from T_0_ measurements in RIF and BDQ experiment. **B.** Relative growth in each strain. Bars denote mean of pooled growth metrics from RIF and BDQ positive controls. **C.** Growth in positive controls of RIF experiment. **D.** Growth in positive controls of BDQ experiment. **E.** Initial ATP measurements by lineage. **F.** Relative growth by lineage. * p < 0.05, *** p < 0.01, p< 0.001 by Kruskal-Wallis with Dunn’s correction for multiple comparisons.

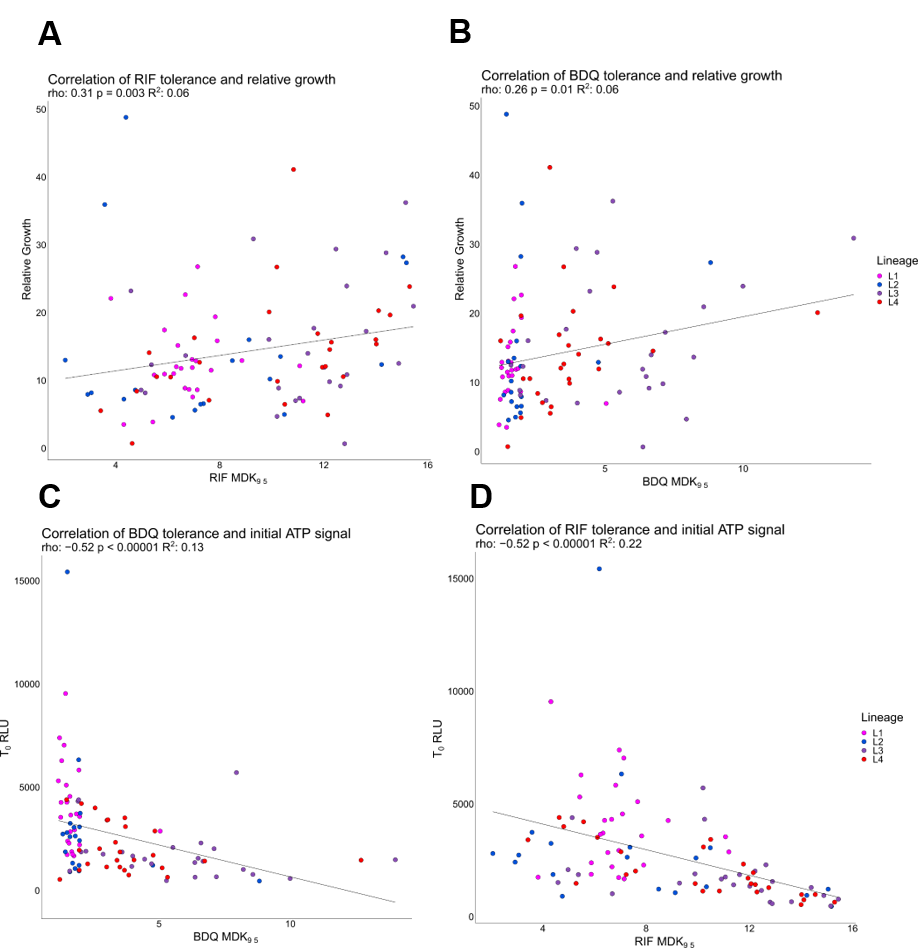

**Figure S10:** Correlation plots relating tolerance to growth and energy phenotypes in strains. **A.** Correlation of RIF tolerance and growth. **B.** Correlation of BDQ tolerance and growth. **C.** Correlation of BDQ tolerance and initial ATP measurement. **D.** Correlation of RIF tolerance and initial ATP measurement. Initial ATP measured values are derived from average of initial reading in both experiments. Growth measurement described in supplementary methods.

**Figure S11:** Phylogeny based representation of strain set and corresponding to MICs for RIF and BDQ. Tips coloured by MTBC lineage L1 = pink L2 = blue L3= purple L4= red. Colour scale depicts MIC.

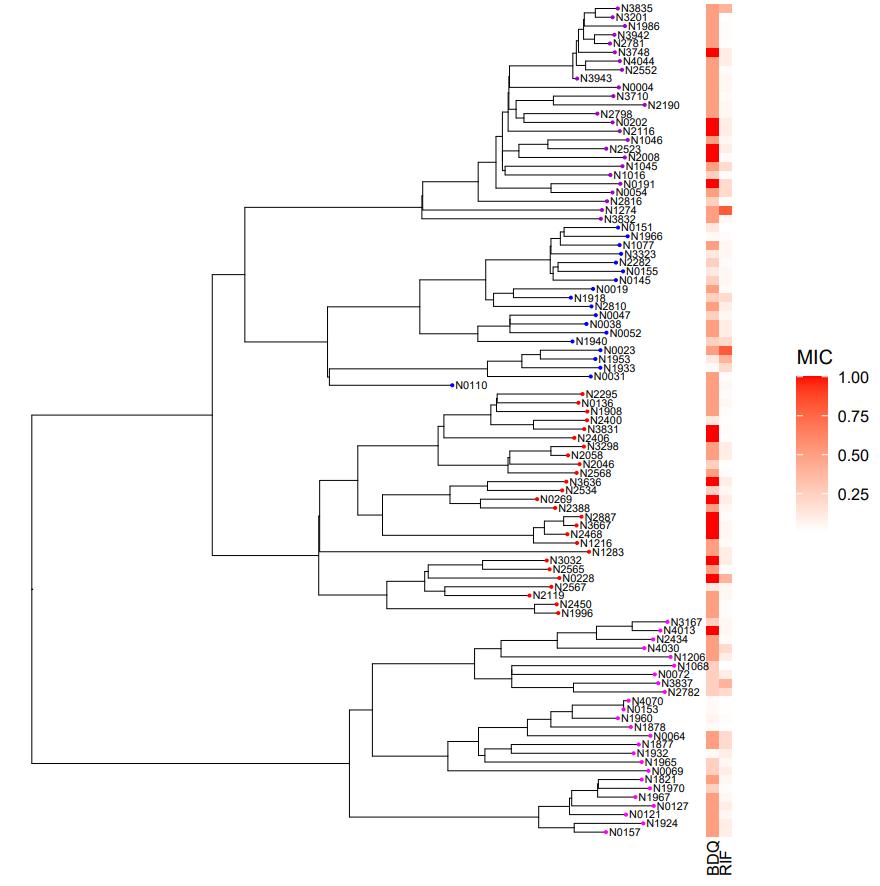

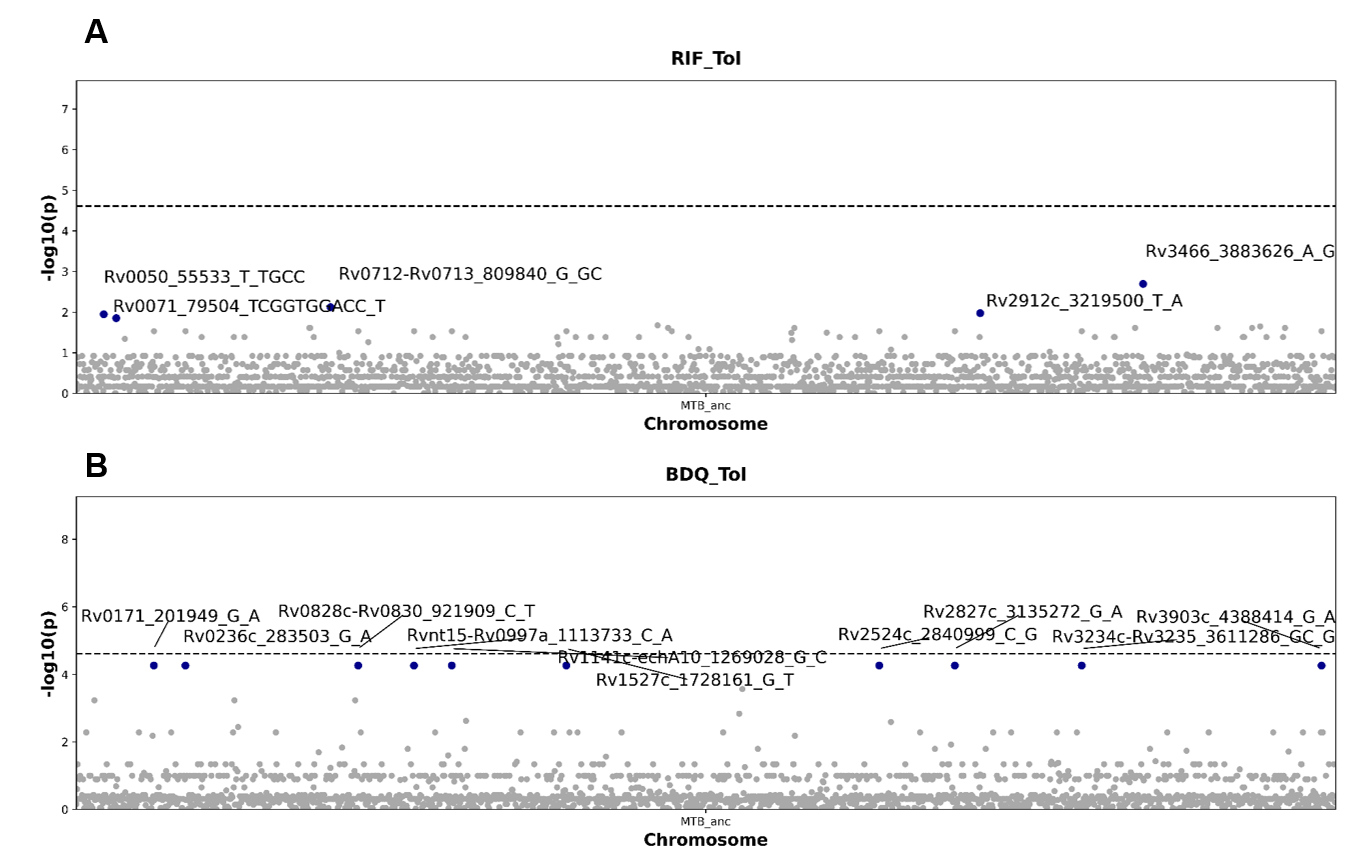

**Figure S12:** GWAS of strain set from RIF (**A**) and BDQ (**B**) rt-TKA experiments. Notable positions highlighted in blue and labelled.

**Table S1**: Sub-threshold, notable genetic polymorphisms associated with high tolerance identified by GWAS

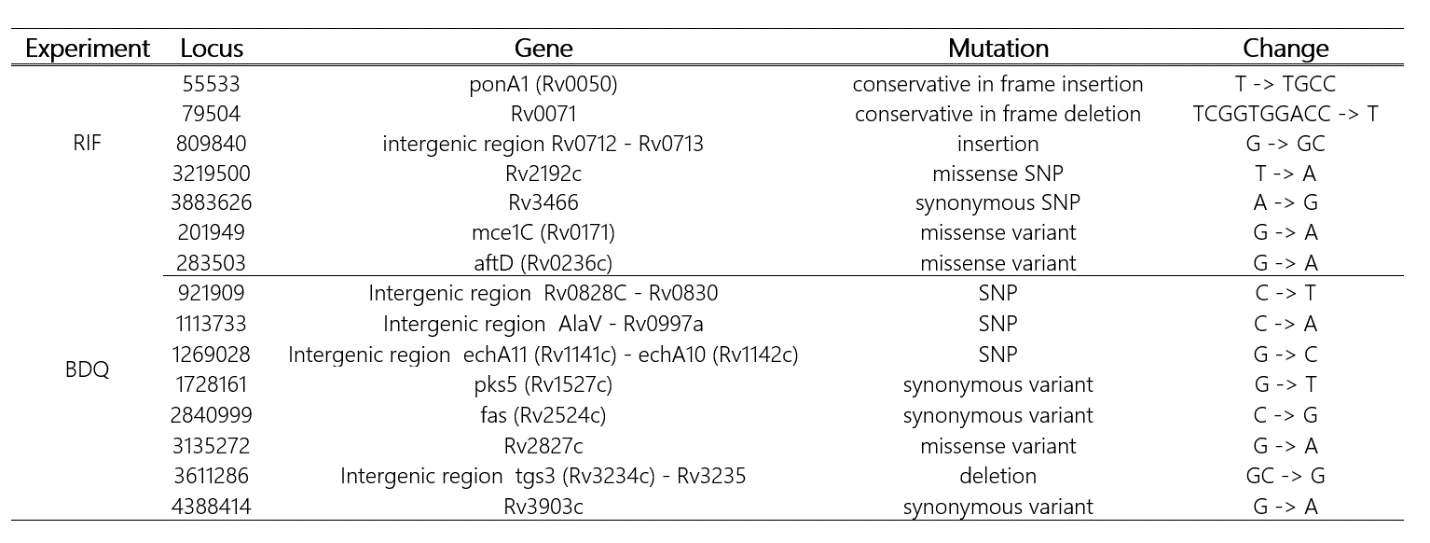

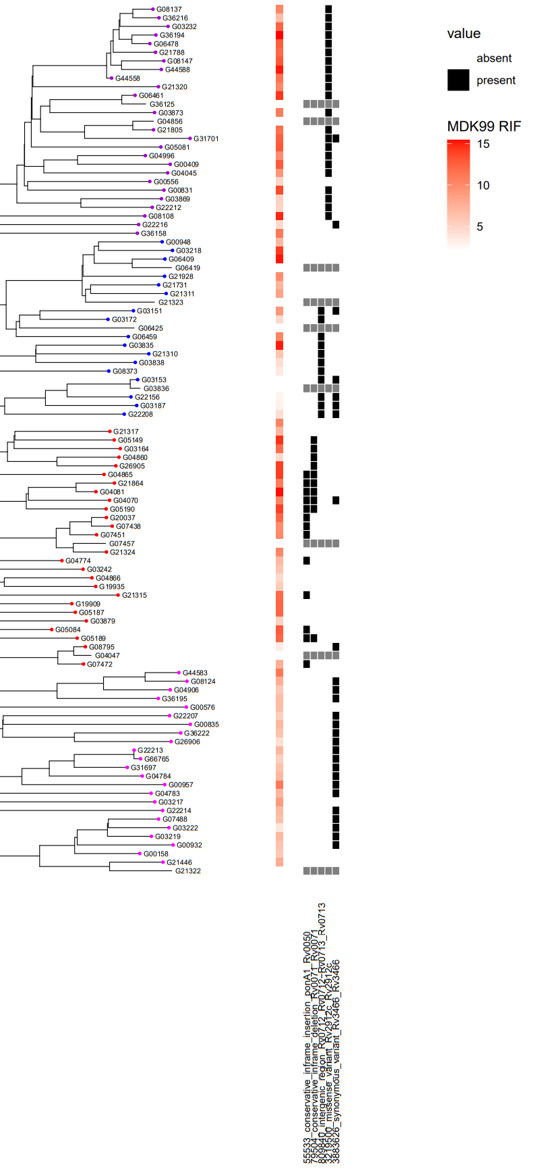

**Figure S13:** Notable positions identified by GWAS of data from RIF experiment. Heat map denotes level of tolerance. Black squares indicates presence, while white space indicates absence. Grey squares indicates genomes that were not assessable.

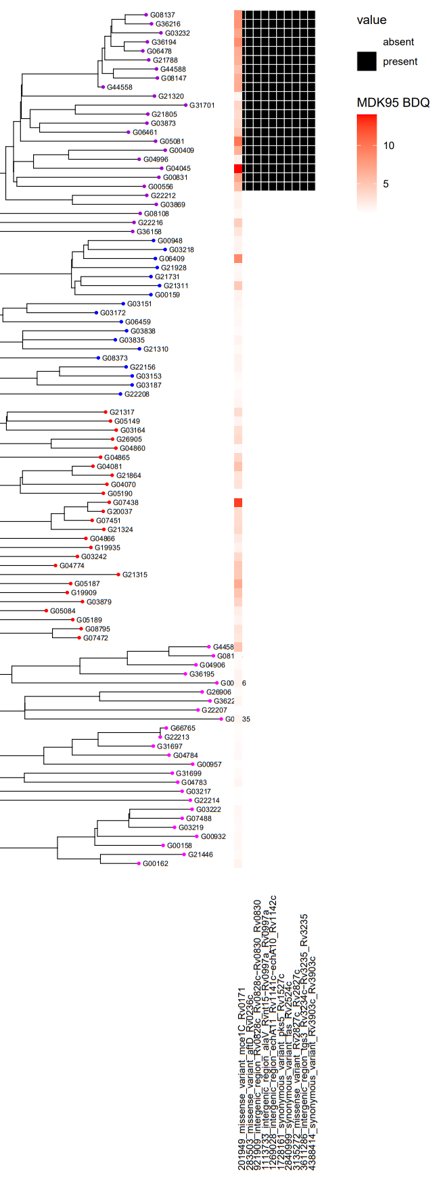

**Figure S14:** Notable positions identified by GWAS of data from BDQ experiment. Heat map denotes level of tolerance. Black squares indicates presence, while white space indicates absence.

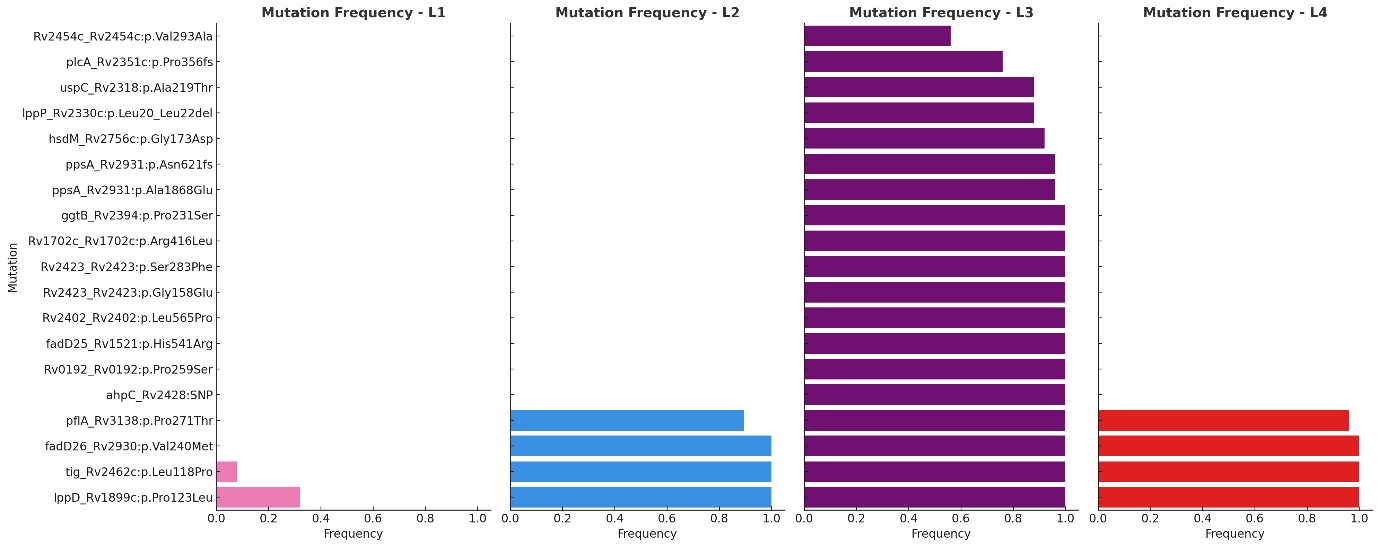

**Figure S15:** Frequency of mutations enriched in high tolerance strains by lineage

**Table S2: Drug response associated positions identified in the literature**

| Gene | Start_coord | End_coord | Reference |
| --- | --- | --- | --- |
| Rv0192_Rv0192 | 223564 | 224664 | <https://doi.org/10.1371/journal.pone.0155127> |
| cobU_Rv0254c | 305825 | 306349 | <https://doi.org/10.1371/journal.pone.0155127> |
| eccA3_Rv0282 | 342130 | 344025 | <https://doi.org/10.1371/journal.pone.0155127> |
| grpE_Rv0351 | 421709 | 422416 | <https://doi.org/10.1371/journal.pone.0155127> |
| prpC_Rv1131 | 1256132 | 1257313 | <https://doi.org/10.1371/journal.pone.0155127> |
| Rv1227c_Rv1227c | 1370292 | 1370825 | <https://doi.org/10.1371/journal.pone.0155127> |
| ppe20_Rv1387 | 1561769 | 1563388 | <https://doi.org/10.1371/journal.pone.0155127> |
| Rv1395_Rv1395 | 1571047 | 1572081 | <https://doi.org/10.1371/journal.pone.0155127> |
| fadD25_Rv1521 | 1712302 | 1714053 | <https://doi.org/10.1371/journal.pone.0155127> |
| ileS_Rv1536 | 1736519 | 1739644 | <https://doi.org/10.1371/journal.pone.0155127> |
| treX_Rv1564c | 1769436 | 1771601 | <https://doi.org/10.1371/journal.pone.0155127> |
| phiRv1_Rv1576c | 1780643 | 1782064 | <https://doi.org/10.1371/journal.pone.0155127> |
| argF_Rv1656 | 1869922 | 1870845 | <https://doi.org/10.1371/journal.pone.0155127> |
| Rv1702c_Rv1702c | 1927211 | 1928575 | <https://doi.org/10.1371/journal.pone.0155127> |
| PE19_Rv1791 | 2029904 | 2030203 | <https://doi.org/10.1371/journal.pone.0155127> |
| cobM_Rv2071c | 2328222 | 2328977 | <https://doi.org/10.1371/journal.pone.0155127> |
| Rv2182c_Rv2182c | 2444586 | 2445329 | <https://doi.org/10.1371/journal.pone.0155127> |
| Rv2184c_Rv2184c | 2445807 | 2446946 | <https://doi.org/10.1371/journal.pone.0155127> |
| mmpS3_Rv2198c | 2462148 | 2463047 | <https://doi.org/10.1371/journal.pone.0155127> |
| gcvT_Rv2211c | 2476042 | 2477181 | <https://doi.org/10.1371/journal.pone.0155127> |
| glnA1_Rv2220 | 2487615 | 2489051 | <https://doi.org/10.1371/journal.pone.0155127> |
| kasA_Rv2245 | 2518115 | 2519365 | <https://doi.org/10.1371/journal.pone.0155127> |
| glpd1_Rv2249c | 2523241 | 2524791 | <https://doi.org/10.1371/journal.pone.0155127> |
| Rv2254c_Rv2254c | 2528520 | 2528975 | <https://doi.org/10.1371/journal.pone.0155127> |
| Rv2264_Rv2263 | 2535641 | 2536594 | <https://doi.org/10.1371/journal.pone.0155127> |
| pitB_Rv2281 | 2553173 | 2554831 | <https://doi.org/10.1371/journal.pone.0155127> |
| cdh_Rv2289 | 2561675 | 2562457 | <https://doi.org/10.1371/journal.pone.0155127> |
| Rv2307c_Rv2307c | 2577851 | 2578696 | <https://doi.org/10.1371/journal.pone.0155127> |
| Rv2308_Rv2308 | 2580419 | 2581135 | <https://doi.org/10.1371/journal.pone.0155127> |
| Rv2323c_Rv2323c | 2595361 | 2596269 | <https://doi.org/10.1371/journal.pone.0155127> |
| Rv2324_Rv2324 | 2596334 | 2596780 | <https://doi.org/10.1371/journal.pone.0155127> |
| Rv2327_Rv2327 | 2599988 | 2600479 | <https://doi.org/10.1371/journal.pone.0155127> |
| recO_Rv2362c | 2643461 | 2644258 | <https://doi.org/10.1371/journal.pone.0155127> |
| phoH1_Rv2368c | 2648916 | 2649974 | <https://doi.org/10.1371/journal.pone.0155127> |
| cfp2_Rv2376c | 2655609 | 2656115 | <https://doi.org/10.1371/journal.pone.0155127> |
| mbtF_Rv2379c | 2657700 | 2662085 | <https://doi.org/10.1371/journal.pone.0155127> |
| mbtB_Rv2383c | 2671593 | 2675837 | <https://doi.org/10.1371/journal.pone.0155127> |
| rpfD_Rv2389c | 2683248 | 2683712 | <https://doi.org/10.1371/journal.pone.0155127> |
| nirA_Rv2391 | 2684679 | 2686370 | <https://doi.org/10.1371/journal.pone.0155127> |
| Rv2402_Rv2402 | 2698529 | 2700457 | <https://doi.org/10.1371/journal.pone.0155127> |
| Rv2423_Rv2423 | 2719597 | 2720643 | <https://doi.org/10.1371/journal.pone.0155127> |
| Rv2425c_Rv2425c | 2721866 | 2723308 | <https://doi.org/10.1371/journal.pone.0155127> |
| ahpC_Rv2428 | 2726193 | 2726780 | <https://doi.org/10.1371/journal.pone.0155127> |
| rplU_Rv2442c | 2740047 | 2740361 | <https://doi.org/10.1371/journal.pone.0155127> |
| Rv2454c_Rv2454c | 2753625 | 2754746 | <https://doi.org/10.1371/journal.pone.0155127> |
| tig_Rv2462c | 2763891 | 2765291 | <https://doi.org/10.1371/journal.pone.0155127> |
| plsB2_Rv2482c | 2786914 | 2789283 | <https://doi.org/10.1371/journal.pone.0155127> |
| Rv2508c_Rv2508c | 2823256 | 2824593 | <https://doi.org/10.1371/journal.pone.0155127> |
| Rv2516c_Rv2516c | 2832710 | 2833513 | <https://doi.org/10.1371/journal.pone.0155127> |
| lppA_Rv2543 | 2866468 | 2867127 | <https://doi.org/10.1371/journal.pone.0155127> |
| secD_Rv2587c | 2914015 | 2915736 | <https://doi.org/10.1371/journal.pone.0155127> |
| fadD26_Rv2930 | 3243697 | 3245448 | <https://doi.org/10.1371/journal.pone.0155127> |
| ppsA_Rv2931 | 3245445 | 3251075 | <https://doi.org/10.1371/journal.pone.0155127> |
| ppsC_Rv2933 | 3255685 | 3262251 | <https://doi.org/10.1371/journal.pone.0155127> |
| ppsD-Rv2934 | 3262248 | 3267731 | <https://doi.org/10.1371/journal.pone.0155127> |
| mas_Rv2940c | 3276380 | 3282715 | <https://doi.org/10.1371/journal.pone.0155127> |
| pks1_Rv2946c | 3291503 | 3296353 | <https://doi.org/10.1371/journal.pone.0155127> |
| fadE30_Rv3560c | 4000432 | 4001589 | <https://doi.org/10.1371/journal.pone.0155127> |
| gid_Rv3919c | 4407528 | 4408202 | <https://doi.org/10.1371/journal.pone.0155127> |
| resR_Rv1830 | 2074841 | 2075518 | [DOI: 10.1126/science.abq2787](https://doi.org/10.1126/science.abq2787) |
| prpR_Rv1129c | 1253074 | 1254534 | <https://doi.org/10.1038/s41564-018-0218-3> |
| ncRv0842_Rv0842 | 938112 | 939404 | Efflux_pump_Miotto_TB_science |
| glpK_Rv3696c | 4138202 | 4139755 | <https://doi.org/10.1073/pnas.1907631116> |
| mamA_Rv3263 | 3643177 | 3644838 | Hypothesised role of Methyl_transferases |
| mamB_Rv2024c | 2268693 | 2270240 | Hypothesised role of Methyl_transferases |
| hsdM_Rv2756c | 3068461 | 3070083 | Hypothesised role of Methyl_transferases |
| tap_Rv1258c | 1406081 | 1407340 | DOI: 10.1093/infdis/jiy710 |
| lppP_Rv2330c | 2603695 | 2604222 | <https://doi.org/10.1038/s41467-021-27616-7> |
| dnaA_Rv0001 | 1 | 1524 | <https://pubmed.ncbi.nlm.nih.gov/33253310> |
| clpX_Rv2457c | 2758208 | 2759488 | [DOI: 10.1126/science.aay3041g](https://science.sciencemag.org/content/367/6474/200.long) |
| purR_Rv3575c | 4017089 | 4018168 | [DOI: 10.1126/science.aay304](https://doi.org/10.1126/science.aay3041) |
| whib7_Rv3197A | 3568401 | 3568679 | <https://doi.org/10.1586/eri.12.90> |
| nrp_Rv0101 | 110001 | 117539 | <https://doi.org/10.7554/eLife.93243.3.sa0> |
| accA2_Rv0973c | 1083747 | 1085750 | <https://doi.org/10.7554/eLife.93243.3.sa0> |
| Rv1319c_Rv1319c | 1480894 | 1482501 | <https://doi.org/10.7554/eLife.93243.3.sa0> |
| cycA_Rv1704c | 1929786 | 1931456 | <https://doi.org/10.7554/eLife.93243.3.sa0> |
| Rv1907c_Rv1907c | 2153235 | 2153882 | <https://doi.org/10.7554/eLife.93243.3.sa0> |
| subI_Rv2400c | 2696644 | 2697714 | <https://doi.org/10.7554/eLife.93243.3.sa0> |
| Rv2893_Rv2893 | 3202420 | 3203397 | <https://doi.org/10.7554/eLife.93243.3.sa0> |
| Rv3483c_Rv3483c | 3902150 | 3902812 | <https://doi.org/10.7554/eLife.93243.3.sa0> |
| proV_Rv3758c | 4203287 | 4204417 | <https://doi.org/10.7554/eLife.93243.3.sa0> |
| Rv3424c_Rv3424c | 3841714 | 3842076 | <https://doi.org/10.7554/eLife.93243.3.sa0> |
| lprA_Rv1270c | 1419014 | 1419748 | <https://doi.org/10.7554/eLife.93243.3.sa0> |
| Rv0792c_Rv0792c | 885837 | 886646 | <https://doi.org/10.7554/eLife.93243.3.sa0> |
| Rv2083_Rv2083 | 2340871 | 2341815 | <https://doi.org/10.7554/eLife.93243.3.sa0> |
| pknH_Rv1266c | 1413960 | 1415840 | <https://doi.org/10.7554/eLife.93243.3.sa0> |
| recN_Rv1696 | 1919683 | 1921446 | <https://doi.org/10.7554/eLife.93243.3.sa0> |
| cut1_Rv1758 | 1989042 | 1989566 | <https://doi.org/10.7554/eLife.93243.3.sa0> |
| lppD_Rv1899c | 2145214 | 2146245 | <https://doi.org/10.7554/eLife.93243.3.sa0> |
| Rv1883c_Rv1883c | 2133231 | 2133692 | <https://doi.org/10.7554/eLife.93243.3.sa0> |
| ctpF_Rv1997 | 2240159 | 2242876 | <https://doi.org/10.7554/eLife.93243.3.sa0> |
| pncA_Rv2043c | 2288681 | 2289241 | <https://doi.org/10.7554/eLife.93243.3.sa0> |
| vapB18_Rv2545 | 2867783 | 2868061 | <https://doi.org/10.7554/eLife.93243.3.sa0> |
| uspC_Rv2318 | 2590518 | 2591840 | <https://doi.org/10.7554/eLife.93243.3.sa0> |
| narK1_Rv2329c | 2601914 | 2603461 | <https://doi.org/10.7554/eLife.93243.3.sa0> |
| plcA_Rv2351c | 2630537 | 2632075 | <https://doi.org/10.7554/eLife.93243.3.sa0> |
| ggtB_Rv2394 | 2688010 | 2689941 | <https://doi.org/10.7554/eLife.93243.3.sa0> |
| cysW_Rv2398c | 2694981 | 2695799 | <https://doi.org/10.7554/eLife.93243.3.sa0> |
| lppB_Rv2544 | 2867124 | 2867786 | <https://doi.org/10.7554/eLife.93243.3.sa0> |
| Rv2488c_Rv2488c | 2797467 | 2800880 | <https://doi.org/10.7554/eLife.93243.3.sa0> |
| Rv2689c_Rv2689c | 3005845 | 3007062 | <https://doi.org/10.7554/eLife.93243.3.sa0> |
| Rv2728c_Rv2728c | 3040766 | 3041461 | <https://doi.org/10.7554/eLife.93243.3.sa0> |
| dinF_Rv2836c | 3142309 | 3143628 | <https://doi.org/10.7554/eLife.93243.3.sa0> |
| pflA_Rv3138 | 3504195 | 3505283 | <https://doi.org/10.7554/eLife.93243.3.sa0> |
| Rv3680_Rv3680 | 4119795 | 4120955 | <https://doi.org/10.7554/eLife.93243.3.sa0> |
| Rv3901c_Rv3901c | 4386365 | 4386814 | <https://doi.org/10.7554/eLife.93243.3.sa0> |
